# Deficient uracil base excision repair leads to persistent dUMP in HIV proviruses during infection of monocytes and macrophages

**DOI:** 10.1101/2020.03.10.985283

**Authors:** Mesfin Meshesha, Alexandre Esadze, Junru Cui, Natela Churgulia, Sushil Kumar Sahu, James T. Stivers

**Affiliations:** Department of Pharmacology and Molecular Sciences, Johns Hopkins University School of Medicine 725 North Wolfe Street Baltimore, MD 21205

## Abstract

Non-dividing cells of the myeloid lineage such as monocytes and macrophages are target cells of HIV that have low dNTP pool concentrations and elevated levels of dUTP, which leads to frequent incorporation of dUMP opposite to A during reverse transcription (“uracilation”). One factor determining the fate of dUMP in proviral DNA is the host cell uracil base excision repair (UBER) system. Here we explore the relative UBER capacity of monocytes (MC) and monocyte-derived macrophages (MDM) and the fate of integrated uracilated viruses in both cell types to understand the implications of viral dUMP on HIV diversification and infectivity. We find that monocytes are almost completely devoid of functional UBER, while macrophages are mainly deficient in the initial enzyme uracil DNA glycosylase (hUNG2). Accordingly, dUMP persists in viral DNA during the lifetime of a MC and can only be removed after differentiation of MC into MDM. Overexpression of human uracil DNA glycosylase in MDM prior to infection resulted in rapid removal of dUMP from HIV cDNA and near complete depletion of dUMP-containing viral copies. This finding establishes that the low hUNG2 expression level in these cells limits UBER but that hUNG2 is restrictive against uracilated viruses. In contrast, overexpression of hUNG2 after viral integration did not accelerate the excision of uracils, suggesting that they may poorly accessible in the context of chromatin. We found that viral DNA molecules with incorporated dUMP contained unique (+) strand transversion mutations that were not observed when dUMP was absent (G→T, T→A, T→G, A→C). These observations and other considerations suggest that dUMP introduces errors predominantly during (-) strand synthesis when the template is RNA. These mutations may arise from the increased mispairing and duplex destabilizing effects of dUMP relative to dTMP during reverse transcription. Overall, the likelihood of producing a functional virus from *in vitro* infection of MC is about 50-fold and 300-fold reduced as compared to MDM and activated T cells. The results implicate viral dUMP incorporation in MC and MDM as a potential viral diversification and restriction pathway during human HIV infection.

## INTRODUCTION

A role for the uracil base excision repair (UBER) pathway in HIV-1 infection of non-dividing macrophage and monocyte immune cells has been of interest for many years in at least two different biological contexts. These contexts arise from two innate immune responses that result in the introduction of dUMP into viral DNA [1–5]. These responses involve either enzymatic cytosine deamination of the viral (-) strand cDNA by APOBEC DNA deaminases [1,2] or the incorporation of dUMP opposite to adenine on either strand of viral DNA during reverse transcription [3,4,6]. The incorporation pathway arises specifically in quiescent cells because [dUTP] is typically higher in such non-dividing cells and most DNA polymerases readily utilize dUTP as a substrate in competition with dTTP [7–9]. The two uracilation pathways are quite unique because cytosine deamination leads to G⍰A transition mutations at specific trinucleotide sequences on (+) strand genomic RNA [10], while dUMP incorporation opposite to adenine occurs on both strands and is not intrinsically mutagenic [3]. Previous studies by us and others indicate that the majority of the dUMP that is present with *in vitro* infected monocyte-derived-macrophages (MDM) arises from dUMP incorporation by reverse transcriptase [4,6]. Once the viral DNA products enter the nuclear compartment, both types of lesions are substrates for uracil excision by the enzyme nuclear uracil DNA glycosylase (hUNG2)[3], the first enzyme in the UBER pathway. Excision by hUNG2 could lead to a variety of different outcomes ranging from viral DNA damage via strand breaks to replacement of dUMP with dTMP, restoring canonical T/A pairs [4,6]. We have previously reported that HIV DNA obtained from alveolar macrophages and circulating blood monocytes from drug naïve and ART patients both contained high levels of dUMP, while T cells from the same patients did not [4]. These findings suggest that uracilation occurs during *in vivo* HIV infection of both macrophages and their monocyte precursors.

It is not clear how HIV-1 remains infective in the hostile deoxynucleotide pool environment of quiescent cells, where overall dNTP concentrations are low and the ratio of [dUTP]/[TTP] is large [11,12]. One plausible explanation is that dUMP/A pairs are well-tolerated in viral cDNA and persist due to very low hUNG2 activity in non-dividing MDM [4]. The intrinsically low activity of hUNG2 in these cells could be further reduced by the ubiquitin-mediated degradation of hUNG2 facilitated by the HIV-1 accessory protein Vpr [4,13–15]. In addition to promoting proteasomal degradation, Vpr has also been suggested to act as a transcriptional repressor of hUNG2 [16]. Another possible outcome in the context of low hUNG2 expression would be slow uracil excision, followed by repair. Over time, and after many excision/incorporation events, proviral dUMP would eventually be replaced with TMP, leading to canonical T/A base pairs. Support for such a replacement mechanism was previously suggested because the levels of dUMP in proviruses decreased over four weeks of culturing infected MDM [4]. The persistence of dUMP in viral DNA and the slow replacement mechanism have implications for viral latency in quiescent cells because *in vitro* and cellular transcription from uracilated DNA templates is greatly reduced even in the complete absence of hUNG activity [9]. This inhibitory effect of dUMP/A pairs arises at the level of transcription factor binding and RNA pol II activity [9,17]. The complexity of this biology that is unique to quiescent cells, challenges efforts to unravel the contributions of the component factors. Nevertheless, the fact that HIV-1 targets the UBER repair pathway at both the transcriptional and protein level indicates an evolutionary pressure to diminish the activity of this pathway during infection [16].

In this study we further explore the nucleotide profiles and UBER activities of MDM and their monocyte precursors with the aim of comparing the outcomes of dUMP incorporation and UBER activity during infection of these cell types (**Fig 1**). The motivation for this work is the previous observation by us and others that HIV DNA is detected in circulating monocytes (MC) and alveolar macrophages (AM) in patients receiving ART who test negative for HIV RNA in blood samples (10-600 HIV copies have been detected per million MC or AM)[4,18,19]. Given that circulating MC are short-lived (*t*_1/2_ ∼ several days)[20], the detection of HIV DNA in these cells of virus-free patients suggests that they arise from contact with latently infected cells in one or more HIV drug-resistant reservoirs, possibly by passing through a compartment that contains infected resident macrophages (or T cells) and then re-entering circulation. Productively infected MC could then seed infection in new tissues (**Fig 1**)[21–23]. We now report that MC are much more deficient in UBER activities compared to MDM such that high levels of viral dUMP persist during the entire lifetime of a circulating MC. When infected MC eventually differentiate into MDM, the existing proviral dUMP can be slowly replaced by TMP using the UBER activities present in MDM. The implications for human HIV infection and potential therapeutic opportunities arising from these findings are discussed.

**Fig 1.**
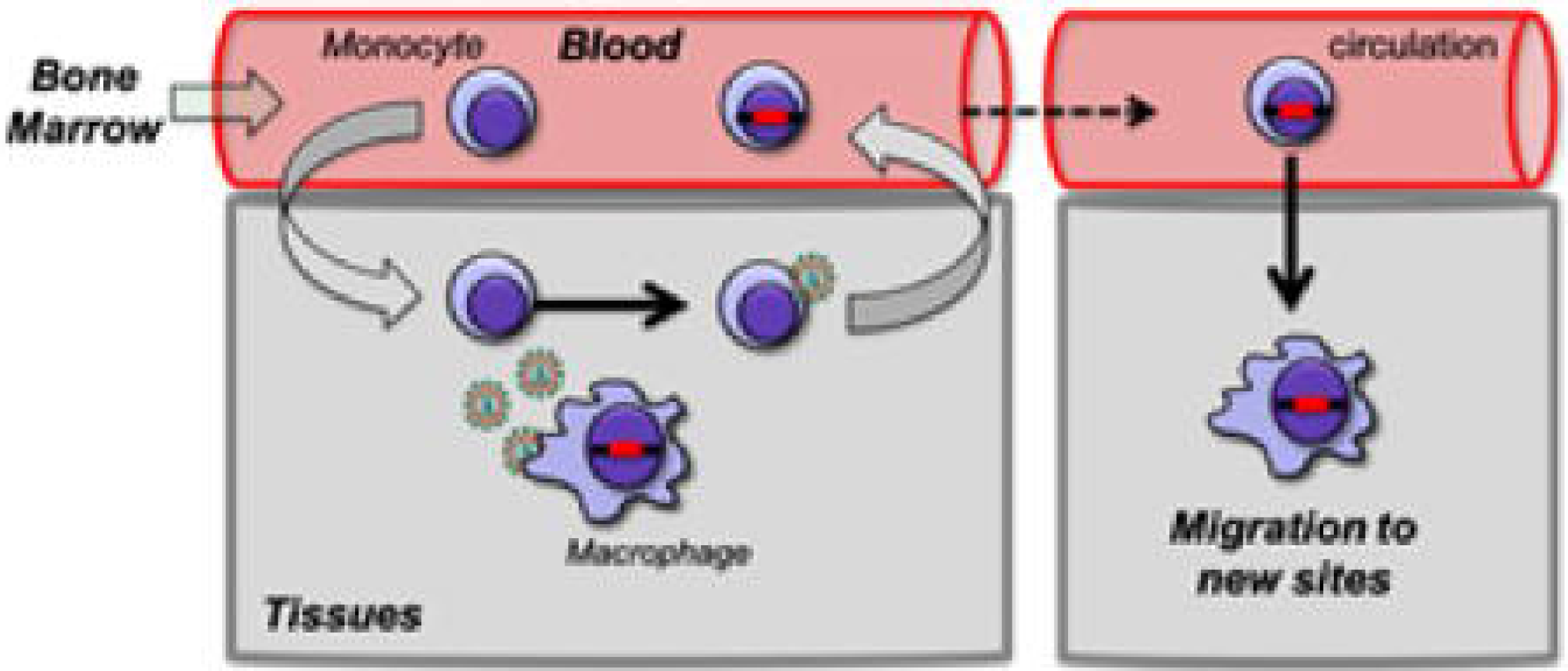
Model for HIV infection of tissue macrophages and infection of new sites via migration of infected monocytes. The model depicts two potential pathways by which macrophages become infected with HIV based on their state of differentiation at the time of virus encounter. In the first scenario, fully differentiated tissue macrophages are infected by contact with virus arising from infected cells passing through the tissue. Alternatively, circulating bone marrow derived monocytes, which are the precursors to some tissue macrophages, could be infected prior to their differentiation into macrophages, either through contact with infected macrophages or T cells. This pathway may be relevant for *in vivo* HIV infection because proviruses have been detected in circulating monocytes of HIV infected individuals who are on ART and have undetectable levels of serum virus.

## Materials and methods

### Cells

Monocytes (MC) were purified from peripheral blood mononuclear cells (PBMC) of HIV negative consenting donors under Johns Hopkins University IRB approval (IRB00038590) using a Ficoll-Hypaque density gradient followed by negative selection using a Pan monocyte isolation kit (Miltenyi Biotech). Monocyte purified by the pan monocyte isolation kit were checked for T-cell contamination by RT-PCR using TCR-β primer pairs which gave 99% purity (see detail in **S1 Supplemental Methods**). In some cases, to prevent cell adherence and differentiation into monocyte-derived macrophages (MDM), MC were cultured up to 7 days in suspension using ultra-low adherence 96-well plates (Corning) using RPMI 1640 (Gibco) supplemented with 10% donor autologous serum, 100 μg/ mL penicillin, 100 μg/mL streptomycin (HyClone), 0.3 mg/ml glutamine and 10 mM HEPES (RPMI-AS). Fully-differentiated MDM were generated by culturing MC under adherent conditions for seven days using RPMI-AS medium supplemented with 10 ng/mL M-CSF (R&D Systems). T cells were purified from PBMC using T cell pan isolation kit (Miltenyi Biotec). T cells were first cultured for three days using stimulating conditions (RPMI supplemented with 10% (vol/vol) donor autologous serum, 100 μg/ mL penicillin, 100 μg/mL streptomycin (HyClone), 0.3 mg/ml glutamine 1 mg/ml of Phytohemagglutinin (Gibco) and, ciprofloxacin (5 mg/ml)). T cells were then infected with HIV^BaL^ and cultured for up to 7 days in RPMI medium additionally supplemented with 10 U/ml recombinant interleukin-2 (Sigma). The dividing cell lines HEK293T (ATCC) and HAP1 (Horizon Discovery) were seeded at 2 million cells per T-75 flask. The culture media for the dividing cells (HI-10) consisted of RPMI-1640 medium + 10% heat inactivated FBS (Sigma), 100 μg/mL penicillin and 100 μg/mL streptomycin. Cells were grown until 70% confluence with media changes every 48 h. The cells were released from the flask using 5 mL of 0.05% trypsin-EDTA (Gibco), followed by washing three times with PBS (Gibco) without CaCl_2_ or MgCl_2_.

### Single nucleotide polymerase extension assay

The extraction of total dNTPs from ∼ 1-2 million cells and quantification of dUTP and dTTP were performed as previously described [3], except that HIV-1 reverse transcriptase (RTase) was used in the extension assay (Millipore-Sigma). Further details are described in the **S1 Supplemental Methods**.

### Western Blotting

Extracts for western blotting were prepared using denaturing conditions with approximately 4 million MC or 2 million MDM that were collected from culture plates by treatment with trypsin. Further details are described in the **S1 Supplemental Methods**.

### Uracil DNA glycosylase activity assay

The hUNG activity present in cell extracts was measured using a molecular beacon fluorescent hairpin DNA substrate containing U/A base pairs as previously described [3,4]. The UNG activity was normalized to the total extract protein. The measured uracil excision activity was fully inhibited by the addition of uracil DNA glycosylase inhibitor protein (UGI) to the extracts, establishing that only hUNG and not any other glycosylase activity was being detected.

### Activity of HIV reverse transcriptase with dUTP and dTTP substrates

To test whether dUMP is an efficient substrate for reverse transcription by HIV RTase, we measured its capacity to distinguish between dTTP and dUTP incorporation using the SNE assay described above. For these measurements we assembled two 100 μL reactions with dNTP mixtures consisting of 1200 fmol dCTP, dGTP, dATP and containing either dTTP or dUTP. The reactions contained 600 fmol of a 5⍰-FAM-labeled DNA template-primer with a single A overhang on the template strand. To these reactions 1 μL (2 units) of recombinant HIV RTase was added and incubated at 37 ⍰C. Ten μL samples were removed at 0, 10, 20, 30, 60, 120, 240, 2400 s, quenched in 40 μL of 98% formamide containing 20 mM EDTA (pH 8.0), and resolved on a 15% denaturing polyacrylamide gel. The initial linear rates of dUTP or dTTP incorporation into the primer strand were measured by fitting data points at < 40% reaction.

### Viruses and infections

The macrophage tropic replicative HIV-1 virus HXB3/BaL was obtained from the NIH AIDS Reagent Program (Catalog #11414) and propagated in MOLT-4/CCR5 cells. Culture supernatants of infected MOLT-4/CCR5 cells were collected at 12 to 15 dpi, centrifuged to clear cellular debris, passed through 0.45 μm filters, aliquoted, and stored at -80 °C. Vesicular stomatitis virus G protein (VSV-G) pseudo-typed HIV-1 virions (HIVNL4.3^(VSVG)^) were generated as previously described (3) by co-transfection of HEK 293T cells with pNL4-3-Δ*E*-eGFP and pVSV-G. pCW57.1.FL.UNG lentiviral particles (LV) were generated by transfecting HEK 293 T cells with; 90 ug of pCW 57.1 hUNG, 90 ug of pMDLg/pRRE packaging plasmid (containing Gag & Pol, Addgene), 36 ug of pRSV-Rev packaging plasmid (containing Rev, Addgene), 18 ug of pMD2.g packaging plasmid (containing VSV-G envelope, Addgene). Transfection was performed using Lipofectamin 2000 (Invitrogen) following the manufacturers protocol. Lentivral particles were then concentrated over 20% sucrose cushion and ultracentrifugation at 28 k RPM for two hours. Viral titer was determined by the ELISA p24 antigen assay (Lenti-X p24 Rapid Titer Kit, TaKaRa). Monocytes were infected immediately after purification by adding HIV-1^BaL^ virus to the culture at MOI between 3 and 5 based on the p24 titer. After infection, monocytes were cultured in two different manners: (i) non-adherent growth conditions in the absence of M-CSF to minimize differentiation by adherence or cytokine mechanisms or, (ii) by adherent growth for seven days in the presence of M-CSF to promote differentiation into MDM over the infection. Infections of fully differentiated MDM were performed after adherent growth in the presence of M-CSF for seven days. Just prior to addition of virus to MDM, the media was replaced with RPMI + 10% dFCS and omitting M-CSF. Monocytes and MDM were also cultured in the presence of 5 mM thymidine (Sigma) overnight to increase the intracellular dTTP levels prior to infection with HIV-1^BaL^ (“thymidine rescue” conditions).

### Viral copy number using qPCR and ddPCR

Viral copy numbers were determined using qPCR. A standard curve for viral copy number quantification was developed using 10-fold serial dilutions of DNA extracted from J-lat cells that contain a single integrated HIV per genome (NIH AIDS Reagent Program). Real time qPCR was performed using a Qiagen Rotor-Gene qPCR instrument using the Rotor-gene Probe PCR kit. Reactions were performed in a 25 ⍰l reaction volume using 0.4 ⍰M forward and reverse primers and 0.2 ⍰M of specific hydrolysis probes. PCR targeted the early or late HIV reverse transcripts (ERT or LRT). Amplification was performed using a two-step program: initial heating at 95 °C for five minutes, followed by 40 cycles of denaturation at 95 °C for 10 seconds and annealing and extension at 60 °C for 30 seconds. For determination of viral copies per cell, simultaneous quantification of the genomic RnaseP (RPP30) gene was performed using published primers and a specific probe as previously described [4]. All primers and probes used in this study are listed in **S1 Table**.

### Determination of HIV proviral DNA using alu-gag nested PCR

To determine copy numbers of HIV proviral DNA, a nested PCR method was used [26]. The first PCR was performed using a forward primer that targeted genomic *alu* sequences randomly located near integrated proviruses (0.2 ⍰M) and an HIV-specific *gag* reverse primer (1.2 ⍰M). Other PCR conditions were 200 ⍰M of each dNTP, 1X LongAmp taq buffer (NEB), 5 units of LongAmp taq DNA polymerase (NEB) and 50 ng of DNA sample extracted from infected cells in a 50 ⍰l final reaction volume. Amplification was performed using the following thermocycler program: initial activation heating 94 °C for two min, followed by 20 cycles of denaturation at 94 °C for 30 sec, annealing at 50 °C for 30 sec and extension at 65 °C for three min and a final extension reaction at 65 °C for ten minutes. The PCR product is diluted 20-fold and five ⍰l of the diluted PCR product is used as an input material for the second PCR reaction, which is performed using LRT forward and reverse primers and probe using the Rotor Gene Probe PCR kit (Qiagen) as described above. Proviral copy numbers were determined using the J-lat cell integration standard as described above. Genomic copy numbers were determined using the RPP30 qPCR measurement described above using the same amount of input total DNA sample used to measure proviral copy numbers.

### Uracil content of viral DNA

Uracil content of viral DNA was determined using either uracil excision droplet digital PCR (Ex-ddPCR) or a similar qPCR method (Ex-qPCR) as previously described [4] with some modifications as described in the **S1 Supplemental Methods**.

### RT-qPCR of extracellular viral RNA

Culture supernatants were collected from infected MC, MDM^EI^ or MDM^LI^ 7-days post infection unless otherwise specified. Supernatants were first spun to remove cellular debris, filtered using 0.22 ⍰m filter and frozen at -80 ^⍰^C until use. RNA was extracted from 140 ⍰l of culture supernatant using the Qiagen mini-Viral Prep kit according to the manufacturer’s protocol. RNA (∼3 µg) was treated with 2 units turbo-DNase (Invitrogen) in 1X turbo-DNase buffer for 30 min at 37 °C, to remove DNA carryover and then purified using Qiagen RNEasy kit. About 1.3 ⍰g of RNA was used to generated cDNA with the Qiagen OmniScript cDNA preparation kit following the manufacturers protocol. Using the cDNA as input material and the Rotor Gene qPCR probe kit (Qiagen), the genomic HIV RNA copies in the supernatants of infected MC and MDM cultures were measured relative to a standard curve developed with the J-lat HIV integration standard cell line (see above). Thermal cycling conditions for qPCR consisted of 95 °C for 5 min, and 40 cycles of denaturation at 95 °C for 10 sec and annealing and extension at 60 °C for 30 sec. The measurements are reported as RNA copies per provirus present.

#### Sequencing of single viral reverse transcripts and genomic RNA copies

Monocytes and MDMs were infected with HIV-1^BaL^ as described above and culture extracts and supernatants were collected at 7 days post infection, filtered and frozen at -80 °C until use. A combination of qPCR and limiting-dilution PCR were performed to obtain samples that contained single copies of viral DNA and RNA as determined by the fraction of qPCR-positive samples at each dilution. Dilutions where one out of five replicates tested positive were taken as clonal according to Poisson statistics [4,9]. Amplified single copies were then sequenced by the Sanger method. Further details are described in **S1 Supplemental Methods**.

### Infectivity of virus isolated from MC and MDM

MOLT-4/CCR5 target cells were infected overnight at an MOI of 0.5 (0.1 pg p24/cell) using HIV-1^BaL^ virus collected from three different producer culture supernatants: (i) MC infected immediately after isolation and cultured for 7 days under non-adherent conditions in the absence of M-CSF to maintain a monocyte-like phenotype, (ii) infected monocytes that were cultured for seven days in the presence of M-CSF [we refer to these cells as “*early infection*” macrophages (MDM^EI^) to indicate that they were infected before differentiating into MDM], and (iii) MDM that were infected after differentiating for 7 days under adherent conditions in the presence of M-CSF [we refer to these infected cells as MDM^LI^) to indicate that infection occurred after the macrophage phenotype was achieved]. One day after the addition of the virus to the MOLT-4/CCR5 cells, the cells were washed and cultured in RPMI 1640 (Invitrogen) supplemented with 10% heat inactivated bovine serum, 100 U/ml penicillin, 100 μg/ml streptomycin, 0.3 mg/ml glutamine and 200 µg/ml neomycin. DNA was extracted at 7 days post infection and proviral DNA levels were measured by alu-gag qPCR. Viral protein 24 (p24) levels in the culture supernatants were measured at day 12 using ELISA.

### Flow Cytometry

Flow cytometric measurements were made on freshly isolated MC or MDM obtained by differentiation for 7 days in the presence of M-CSF. Prior to flow cytometry, adherent MDM were washed twice with Hank’s balanced salt solution (HBSS) and released from the culture flask by treating with Accutase (Cell Technologies) for 30⍰min at 37 °C. MC and MDM were resuspended at a concentration of 10^6^ cells/mL in 1 × HBSS (pH 7.2), 5⍰mM EDTA and 0.5% BSA and passed through a 35⍰⍰m nylon mesh (BD Biosciences) before injection on a FACSCalibur flow cytometer (Becton Dickinson). For each cell type, 20,000 events were recorded for analysis. Dot plots were created using forward scatter (FSC-H) and side scatter (SSC-H) measurements to determine the size and granularity differences. All data were analyzed using Cell Quest software.

### Light Microscopy

MC were prepared as described above using the Pan-Monocyte isolation kit from Miltenyi. The obtained MC were resuspended in fresh RPMI medium and allowed to adhere on a glass slide for an hour before collecting images at 20x magnification. MDM were differentiated from MCs as described above using RPMI media containing 10% autologous sera and M-CSF. After seven days, the adherent cells were visualized using 20x magnification and images were collected. Images were converted to 8-bit grayscale using ImageJ software.

#### Statistics

The data uncertainties are expressed as the mean ± 1 SD. Unless otherwise indicated in the figure legends, data are derived from at least three independent experiments using cells isolated from at least three donors. Experiments using cells from individual donors were performed in triplicate. Data were analyzed using the Prism 8.0 statistical program from GraphPad Software. Pairwise statistical analysis of test groups was performed by a two-tailed unpaired Student’s *t*-test, assuming equal variances between groups. The significance of the mutation frequencies observed between two groups was determined using a chi-squared analysis.

## RESULTS

### dUTP/TTP in MDM and MC is high but variable with blood donors

In previous studies dUTP/dTTP ratios in the range 20:1 to 60:1 were reported [4,12,24], while the ratio was essentially zero in dividing cells due to the expression of dUTPase in replicating cells [4]. These different estimates of intracellular dUTP/dTTP may arise from the combined challenges of measuring low levels of dNTPs present in non-dividing cells, the inherent difficulty in accurately determining the ratio when one of the nucleotides is present at an extremely low level, intrinsic differences in dNTP pools in different donors, or differences in culture conditions or measurement methods.

We first extended the dTTP and dUTP measurements to freshly isolated monocytes with the expectation that the levels in freshly isolated cells would most closely match *in vivo* levels. A single nucleotide polymerase extension (SNE) assay was used where a 22 mer DNA primer-template containing a single adenine overhang on the template strand was extended by the dUTP or dTTP present in cell extracts (**Fig 2a**)[3]. The assay takes advantage of the enzyme dUTPase to remove dUTP in the extract and allow measurement of dTTP in isolation (see methods). For these assessments, the [dTTP + dUTP] present in MC and MDM extracts were compared with the Hap1 dividing cell line (**Fig 2b** and **S1 Fig**). We found similarly low levels of [dTTP + dUTP] in freshly isolated MC extracts as compared to MDM that had been differentiated over seven days in the presence of M-CSF (1.1 to 2.5 pmol dU(T)TP/million cells). For comparison, the combined [dTTP + dUTP] level in the Hap1 cell line is 25 to 70-fold higher (∼70 pmol dU(T)TP/million cells). Depleting dUTP in these dNTP extracts by the addition of dUTPase allowed calculation of dUTP/dTTP in each cell type (**Fig 2c**). Although freshly isolated MC and differentiated MDM showed similar elevated dUTP/dTTP ratios in the range 1:4 to 1:1 depending on the donor, the ratio was essentially zero for the HAP1 cells. Based on these and previous studies, we conclude that significantly elevated dUTP/dTTP is an intrinsic aspect of MC and MDM metabolism that differs from dividing cells.

**Fig 2.**
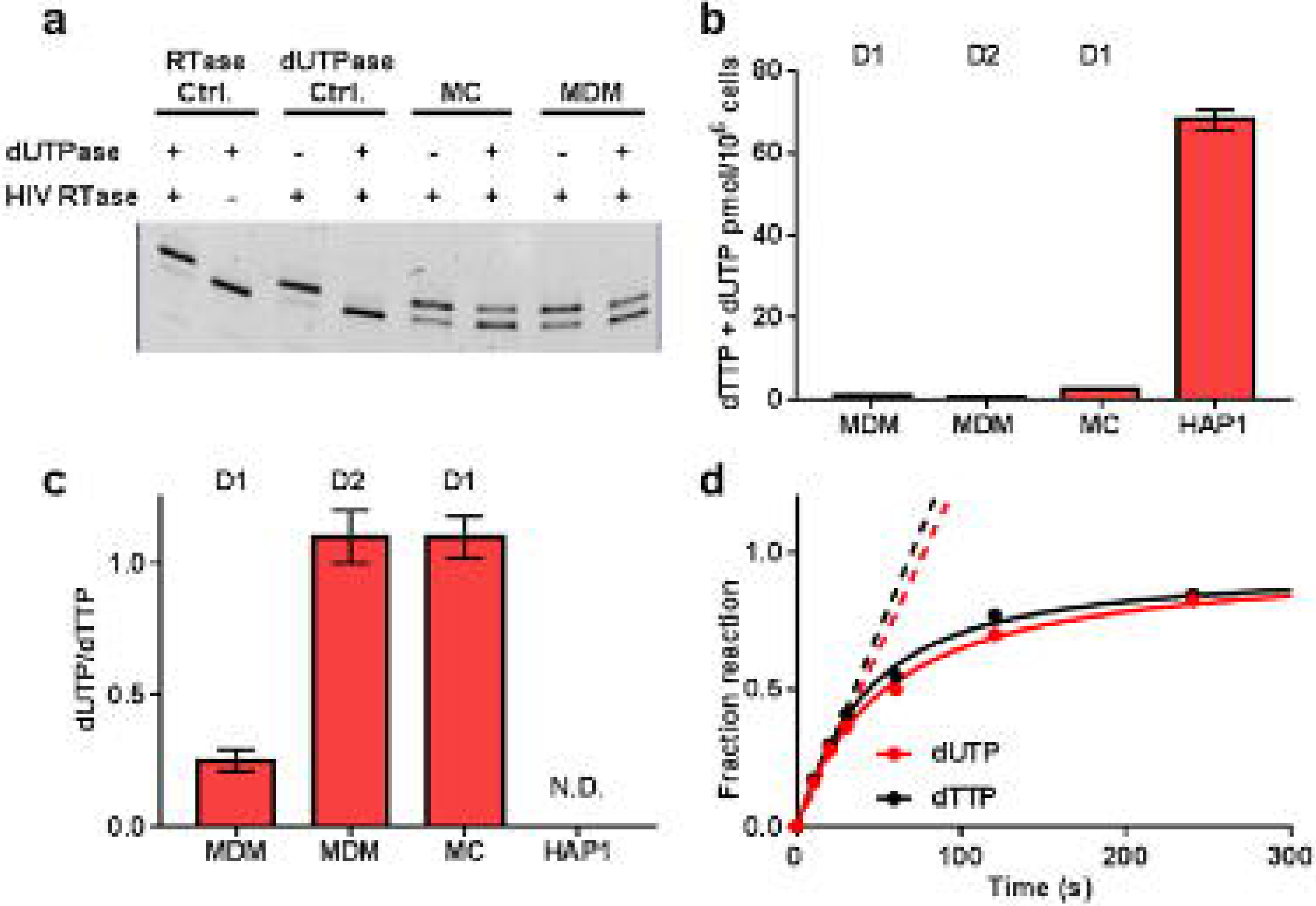
dUTP and dTTP pool measurements in extracts from MDM, MC and comparison with Hap1 dividing cells. A single nucleotide extension assay (SNE) was used to measure dUTP and dTTP levels in MDM, MC and the Hap1 dividing cell line (Fig. S1). The specific measurement of dUTP in a mixed pool of TTP + dUTP is accomplished by the *in vitro* conversion of dUTP dUMP + PP_i_ using dUTPase prior to the SNE assay. (**a**) Typical SNE assay to determine dUTP levels in cell dNTP extracts. The RTase control (dTTP only) establishes that RTase is required for converting the *n* bp substrate into *n + 1* and that dUTPase does not inhibit extension in the absence of dUTP (see methods). The dUTPase control establishes that RTase can fully extend the probe by a single nucleotide in the presence of a dNTP mixture with dUTP replacing dTTP. The addition of dUTPase completely abolishes extension under these conditions. (**b**) Combined dTTP + dUTP pool measurements for MDM and MC from two donors (D1 and D2) and the HAP1 dividing cell line. Error bars are standard errors from triplicate measurements. (**c**) dUTP/dTTP ratio for the various cell types. ND, not detected. Error bars are standard errors from triplicate measurements. (**d**) Determination of the relative activity of HIV RTase for incorporation of dTTP or dUTP opposite to adenine. The solid line is a theoretical fit to the entire time course, while the dashed lines are initial rate linear fits, which are statistically indistinguishable. The concentration of the template DNA was 50 nM and concentration of each dNTP was 100 nM, which is the estimated concentration of these dNTPs in MDM or MC.

Although a range of dUTP/dTTP values have been reported for MDM [4,24], all of the measurements are consistent with low dNTP pools and significant levels of dUTP for both MC and MDM. The current measurements indicate that uracilated viral cDNA will be produced during infection of both MC and MDM and that the UBER capacity of these cells could impact the outcome of infection. However, unlike previous studies where dUTP/dTTP > 20, the more or less balanced ratio (∼1) indicates that viral dUMP residues have the potential for being replaced by dTMP after multiple repair cycles (i.e. each replacement attempt has a 50:50 chance of replacing dUMP with dTMP, but eventually all dUMP would be repaired).

To evaluate whether HIV RTase can discriminate between dUTP and dTTP, and therefore bias dUMP incorporation away from the level expected from the dUTP/dTTP ratio, we measured the RTase activity *in vitro* using both dUTP and dTTP as substrates (**Fig 2d**). Using a low dNTP concentration that approximated that calculated for non-dividing cells, we were unable to detect any selectivity of RTase for either nucleotide. Thus, the relative amounts of dUMP and dTMP in HIV reverse transcripts should reflect the cellular dUTP/dTTP.

### MC and MDM have different uracil base excision repair (UBER) activities

Given that RTase shows no discrimination between dUTP and dTTP, the observed ratios of dUTP/dTTP of 0.25 to 1 for MDM and MC indicate that 99.9 % of HIV DNA products would have at least one dUMP incorporation for every turn of the DNA helix assuming a random sequence containing 50% A/T base pairs. These putative densely spaced uracils would be subject to excision by the UBER pathway. Accordingly, we were interested in the relative levels of seven enzymes involved in deoxyuridine metabolism in MC and MDM as compared to the Hap1 dividing cell line (**Fig 3**). These enzymes included uracil DNA glycosylase (hUNG), AP endonuclease 1 (APE1), DNA polymerase β (pol β), ligase III⍰ (the ligase isoform expressed in non-dividing cells)[25], the dNTPase sterile alpha motif histidine-aspartate domain protein 1 (SAMHD1), and the DNA cytidine deaminases APOBEC3A (A3A) and APOBEC3G (A3G). The western blots and activity measurements for extracts collected from uninfected cells and MC and MDM that were infected with the CCR5 tropic HIV-1^BaL^ viral strain revealed the following general trends (**Fig 3a-3i**) as summarized graphically in **Fig 3m-3s**. First, MC and MDM show similarly low levels of expression of the first enzyme in the UBER pathway (hUNG), which are about 25 to 50-fold lower than the Hap1 reference line. These low hUNG activity levels, which could result from either the mitochondrial (hUNG1) or nuclear isoforms (UNG2), indicate that viral uracils may not be efficiently excised once viral DNA enters the nuclear compartment. Surprisingly, the activity of the next UBER enzyme, APE1, is highly expressed in both MDM and Hap1 cells, indicating that the rate-limiting step for repair of uracil in non-dividing cells is likely the initial excision event by hUNG2. The next two enzymes in the pathway, pol β and lig III⍰, are almost undetectable in MC but much more prevalent in MDM, although not to the same level as the Hap1 line. This important distinction between MC and MDM suggests that uracilated viruses generated by direct infection of MC cannot be repaired until the MC differentiate into MDM. As expected, SAMHD1 dNTPase is highly expressed in both MC and MDM, but not Hap1 cells, consistent with the different dNTP pool levels present in these cell types (**Fig 2b**). Both deaminase enzymes, A3G and A3A, are highly expressed in MC, but not MDM or Hap1 cells. Despite the high expression levels of A3G and A3A, the high dUMP content of viral DNA in MC is not derived from intrinsic A3A or A3G activity—at least not with HIV-1^BaL^ virus that encodes viral infectivity factor (*vif*)(see below). None of the expression data were noticeably affected by whether the extracts were prepared before or after BaL virus infection (**Fig 3**). We performed additional quantitative RT-qPCR measurements of mRNA expression levels of these genes which were fully consistent with the above trends (**S2 Fig**).

**Fig 3.**
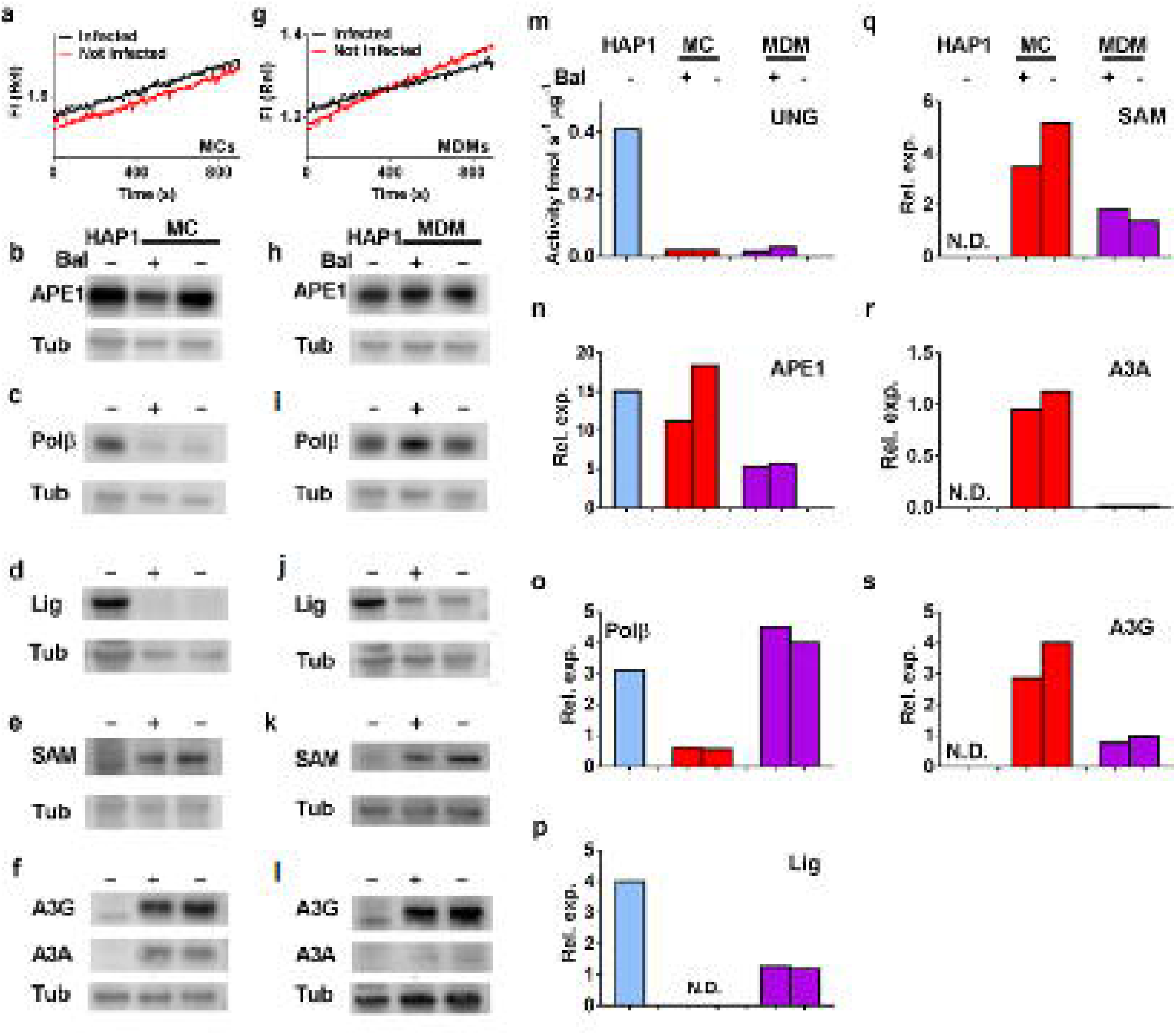
Determination of uracil base excitation repair (UBER) capacity of infected and uninfected MCs, MDMs and comparison with the HAP1 dividing cell line. MC or MDM were infected with replication competent CCR5 tropic HIV-1^BAL^ virus at moi = 5. Extracts from MC (2^nd^ lane in panels **b, c, d, e, f**) and MDM (3^rd^ lane in panels **h, I, j, k, l**) using uninfected (-) or infected cells (+) at 3 days post-infection were processed for western blotting. Blots were performed using 10 μg of total cell protein, except for Lig IIIα, where 20 μg was used. Due to the low levels of hUNG present in non-dividing cells, native extracts (10 μg) were prepared for measurement of UNG enzymatic activity using a sensitive real-time fluorescence assay (see methods). The specific measurement of UNG activity was established by the addition of UNG inhibitor UGI (4 μM), which completely inhibited the measured activity. Panels **a** to **i** show western blots or activity assays for detection of (**a, g)** uracil DNA glycosylase (UNG); (**b, h**) AP endonuclease 1, (APE1); DNA polymerase β, (polβ) (**c, i)**; (**d, j**) ligase III (Lig); (**e, k)** SAMHD1 (SAM); and (**f, l)**. APOBECA3A (A3A) and APOBECA3G (A3G); tubulin (Tub) loading control was used in each blot. Panels **m** to **s** show the relative expression level (Rel. exp.) or activity of each enzyme for each cell line as indicated in each panel. Relative expression was calculated by dividing the band intensity of the protein of interest by the intensity of loading control after correction for background.

### HIV Infection of MC and MDM at early and late stages of differentiation

To explore the origins and possible fate of infected circulating monocytes observed in HIV patients on ART, we performed HIV infections using freshly isolated, undifferentiated MC rather than fully differentiated MDM. We began this exploration by measuring the levels of dUMP present in proviral DNA under three distinct conditions (i) MC that were infected with HIV-1^BaL^ immediately after isolation and then maintained as undifferentiated monocytes for 7 days by culturing under nonadherent conditions in the absence of M-CSF, (ii) MDM infected at the monocyte stage and then allowed to differentiate in the presence of M-CSF (*E*arly *I*nfection, MDM^EI^) and, (iii) MDM that were infected after seven days of differentiation in M-CSF (*L*ate *I*nfection, MDM^LI^) (**Fig 4a**). For comparison, we infected activated T cells using the same virus stock and MOI. Proviral dUMP levels were determined using the alu-gag Ex-qPCR experiment after isolating genomic DNA at 1, 3, 7, 14- and 28-days post infection (dpi). To prevent multiple round infection, 0.2 µM enfuvirtide (T-20) was added one day after the initial infection (**Fig 4a**). The Ex-qPCR analysis determines the fraction of integrated HIV viruses in a DNA sample that contain one or more dUMP residues on each DNA strand of a DNA amplicon that contains a 650 bp region of the 5’ LTR and a 700 bp portion of the gag gene [4,26]. For MC, which were cultured using non-adherent conditions in the absence of M-CSF to maintain a monocyte-like phenotype (**S3 Fig**), no proviral DNA was detected at 1 dpi, but by day three nearly 100% of the proviral copies contained dUMP, which persisted until the end of the experiment (7 dpi)(**Fig 4b**). Due to limitations in the length of time MC can be maintained using nonadherent conditions, this experiment could not be continued beyond seven days. For MDM^EI^, which were infected immediately after isolation of MC and then cultured using adherent conditions in the presence of M-CSF to immediately begin their differentiation into MDM, over 50% of the provirus contained dUMP at 1 dpi, which increased to almost 100% at 3 dpi. Unlike MC grown under non-adherent conditions in the absence of M-CSF, the copies of HIV in MDM^EI^ that contained dUMP decreased by 7 dpi (Frac U ∼ 75%) and even further at 14 dpi (Frac U ∼ 20%). For MDM^LI^, which were infected after complete differentiation, the proviral copies that contained dUMP were initially lower than MC or MDM^EI^ (Frac U ∼ 15% at 1 dpi and 60% at 7 dpi), but ended up at the same level as MDM^EI^ at 14 dpi (Frac U ∼ 20%). We attribute the lower Frac U values of differentiated MDM at early times after infection to the greater rate of reverse transcription in a small sub-population of permissive MDM (<10%) which have higher dNTP pools and low dUTP [4]. As the post-infection time increases, the slower replicating uracilated DNA products in the major MDM population increasingly contributes to the bulk measurement. The control T cells showed no viral-associated dUMP at any time during infection, consistent with our previous finding (4).

**Fig 4.**
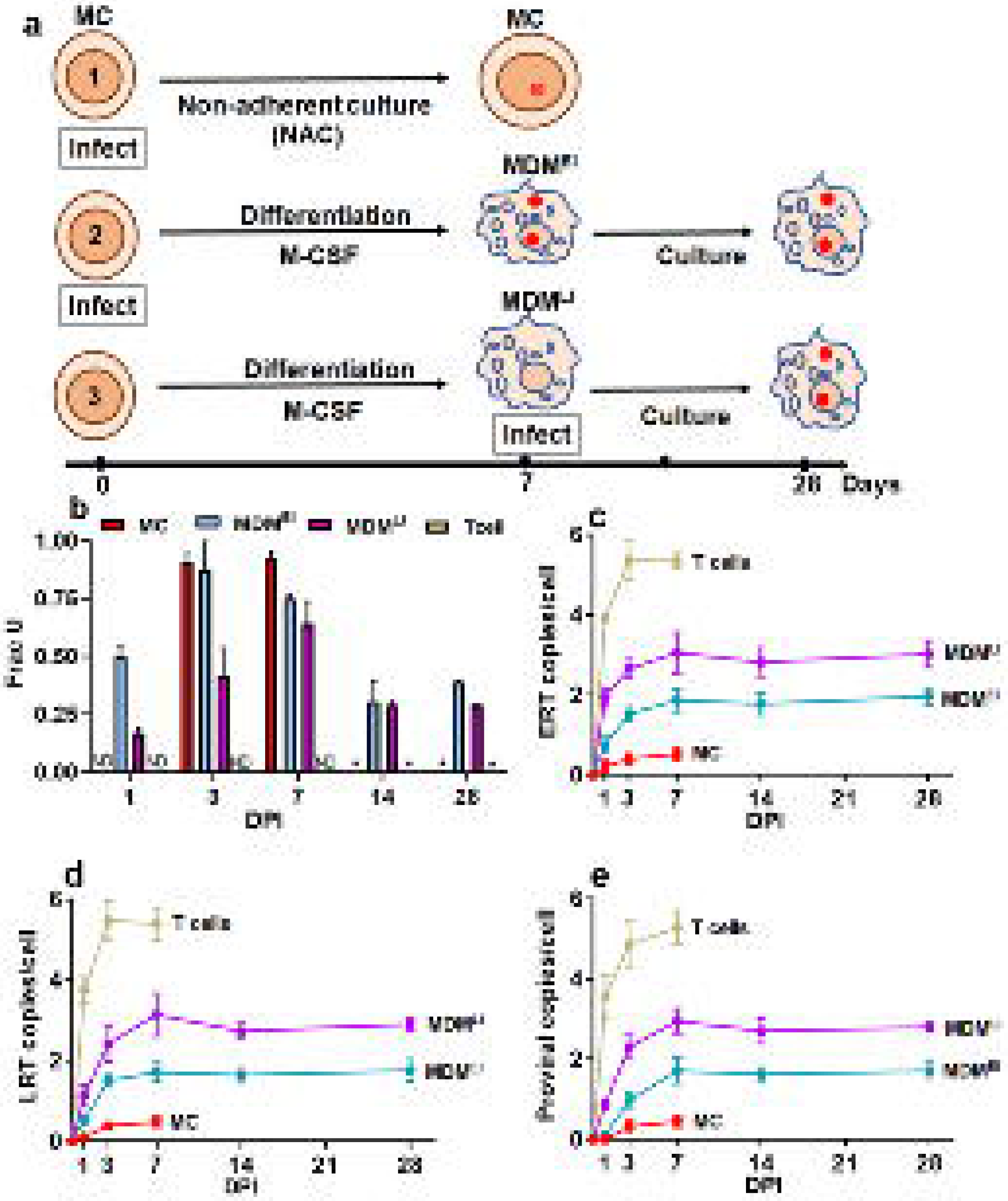
Characteristics of HIV infection in MC and macrophages infected before differentiation (MDM^EI^) and after differentiation (MDM^LI^). (**a**) Infection of MC and MDM with HIV-1^BaL^ virus was performed using three conditions (moi = 5). In one condition, MC were immediately infected and then cultured using nonadherent conditions without M-CSF for seven days to maintain a monocyte phenotype. MDM^EI^ refers to “early infection” of freshly isolated MC, which were subsequently differentiated into MDM under adherent conditions in the presence of M-CSF. MDM^LI^ refers to “late infection” where MDM were first fully differentiated before infection. In all cases, HIV fusion inhibitor drug, enfuvirtide, (0.2 µM final concentration) was added to culture medium after 24 hours of infection to prevent multiple rounds of infection. (**b**) Fraction of provirus in MC, MDM^EI^ and MDM^LI^ that contain uracil as determined by alu-gag Ex-qPCR. ND: not detected (**c**) Time course for appearance of early reverse transcription (ERT) products as determined by RT-qPCR using a primer set targeting the 5’LTR region of the HIV genome. (**d**) Time course for appearance of late reverse transcription (LRT) products as determined by RT-qPCR using a primer set targeting the LTR region of the HIV genome. (**e**) Time course for appearance of proviral DNA copies determined by alu-gag qPCR. Identical infections activated T cells were used as controls and are shown in panels **d, e** and **f**. Abbrev: ND, not detected. * not done. All data are averages obtained from three blood donors.

We then investigated the kinetics for appearance of early and late reverse transcripts (ERT, LRT) and proviral DNA using the three infection conditions. The kinetics for forming ERT and LRT products followed the trend MDM^LI^ > MDM^EI^ ≫ MC (**Fig 4c, d**), with ERT and LRT copy numbers for MDM^EI^ and MC about 2 and 6-fold lower at 7 dpi as compared to MDM^LI^. A similar trend was observed for the proviral copy numbers (**Fig 4e**). The control infections of activated T cells showed both faster reverse transcription kinetics (∼70% complete in ∼1 day, **Fig 4c, d, e**) and a greater number of integrated proviruses.

### dUMP in proviral DNA does not arise from cytidine deaminase activity in MC

Given the presence of APOBEC enzymes in MC, and to a lesser extent MDM as judged by immunoblotting (**Fig 3a, g**), we wanted to confirm that most of the dUMP in HIV proviral DNA was derived from incorporation of dUMP by RTase. One established way to test this is to add high levels of thymidine (dThyd) to the cell culture media prior to infection to increase the levels of intracellular TTP and then look for a reduction in the number of viral DNA products that contain dUMP [4]. We infected both MC and fully differentiated MDM in the presence of 5 mM dThyd and measured the copies of uracilated proviruses at 7 dpi using the Alu-gag Ex-qPCR method (**Fig 5**). For both cell types we observed a 4-8-fold reduction in uracilated viral copies, indicating a predominant role for dUMP incorporation. A further reduction in uracilated viral copies is not expected due to the minor population of highly permissive MDM that do not contain high dUTP levels [4,27]. A minor role for APOBEC DNA cytidine deamination is confirmed by our proviral DNA sequencing results reported below, which show a low frequency of G⍰A transition mutations on the proviral (+) strand DNA at known APOBEC hotspots.

**Fig 5.**
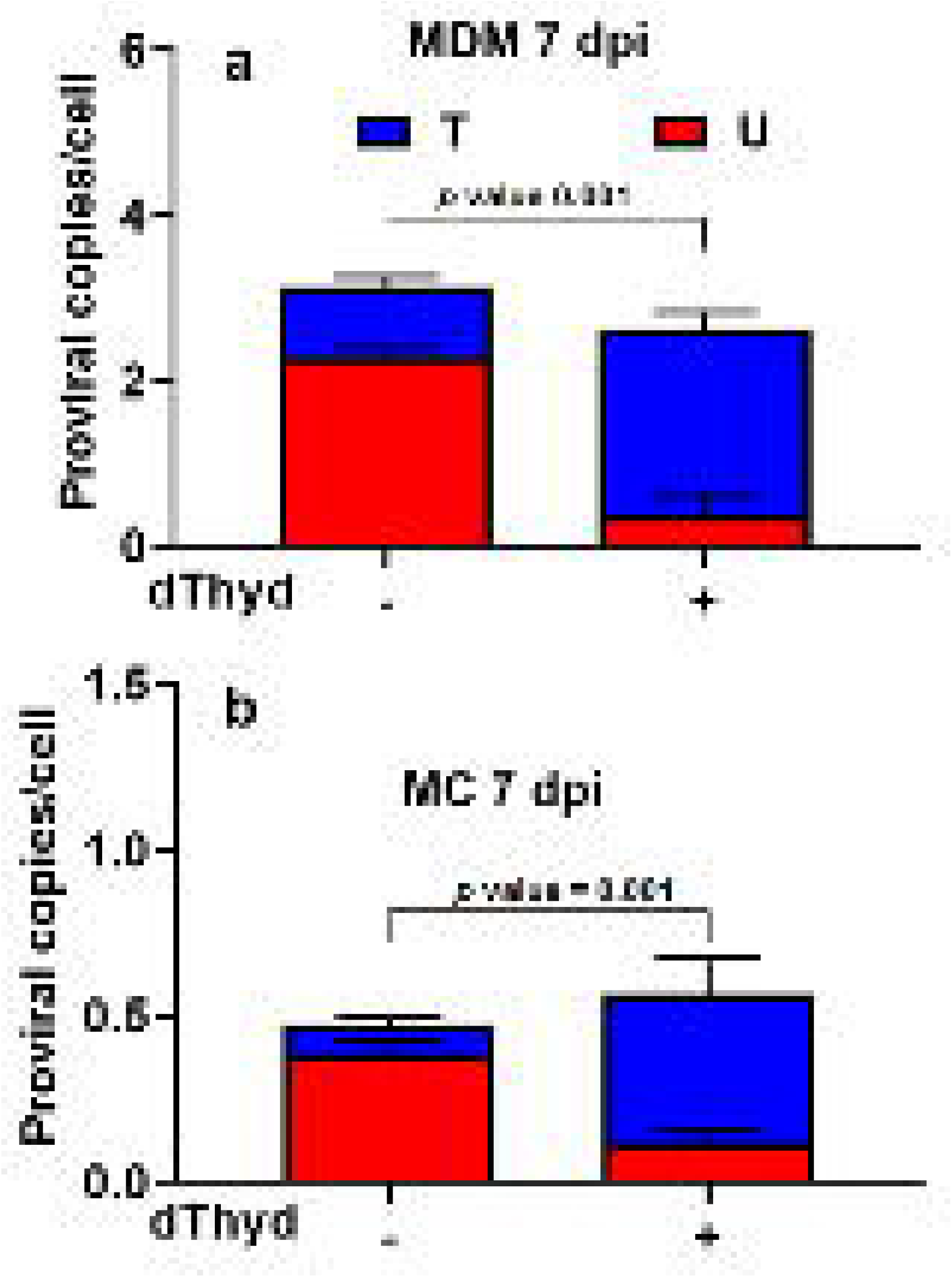
dUMP in HIV DNA during infection of MC and MDM arises predominantly from dUTP. MC and MDM were infected with HIV-1^Bal^ virus at an MOI of 5 (1 pg p24/cell) in the presence (+) and absence (-) of 5 mM deoxythymidine (dThyd). Copy number measurements of uracilated (red) and non-uracilated (blue) proviruses were made 7 dpi using the Alu-gag Ex-qPCR method. (**a**) MDM and (**b**) MC infected in the absence and presence of dThyd. *p*-values from an unpaired Student’s *t*-test are shown.

### Effect of viral dUMP on proviral DNA and extracellular RNA sequences

Although dUMP incorporation is not expected to introduce mutations because its Watson-Crick hydrogen bond donor and acceptor groups are identical with thymidine, dUMP/A base pairs have reduced thermodynamic stability in the context of B DNA [28], increased base pair dynamics [29], and dUMP has an increased propensity to form dUMP/G wobble mismatches relative to thymine due to its larger keto-enol tautomerization constant. The reduction in duplex stability arising from dUMP has been attributed to the lower electron density of the pyrimidine ring system of uracil relative to thymidine, which weakens base stacking [29]. Thus, we were interested if any of these potential effects of dUMP incorporation could be detected in cDNA sequences produced from reverse transcriptase or viral RNA genomes produced by RNA pol II transcription.

To investigate proviral DNA sequences produced by reverse transcription in both MC and fully differentiated MDM, we used HIV-1^BaL^ virus to infect freshly isolated MC cultured under non-adherent conditions and also MDM after 7 days of culturing. Both cell types were cultured for seven days after infection and total DNA was isolated and diluted to the single copy level. Single proviral clones were then amplified using ES7 and ES8 env primers (see **S1 Table**). Wells that were positive for HIV DNA clones by qPCR were reamplified using primers that generated a 592 bp amplicon covering the V3 and V4 regions of *env*, followed by sequencing using the Sanger method. The viral sequences were compared with the lab reference sequence to determine the mutation frequencies and types. For proviral sequences isolated from MC and MDM, the viral mutation frequency (∼1.5 × 10^−3^) and mutational spectrum were similar (**Table 1, Fig 6a, b).** For both cell types, 26 to 38% of the isolated proviral clones contained substitution mutations, with the majority appearing as transition mutations (∼70-80%), and the remaining being more unusual transversion mutations (20% to 30%). Two (+) strand G⍰A mutations detected in both MDM and MC infections might be attributed to enzymatic cytosine deamination on the viral (-) strand cDNA based on the sequence preferences of A3A or A3G (**S2 Table**). However, for MDM we cannot exclude that these apparent enzyme derived mutations arose from chance misincorporation because four G→A mutations occurred on the viral (-) strand cDNA (corresponding to C→T on the positive strand). These mutations in MDM cannot be attributed to APOBEC activity because of their sequence context and occurrence on the (-) strand (**S2 Table**). The two (+) strand G→A mutations observed in the MC derived samples could arise from the high A3A activity in these cells. Finally, 40% and 55% of the proviral mutations in MDM and MC led to codon changes and could therefore affect viral fitness (**S2, S3 Table**).

**Table 1.**
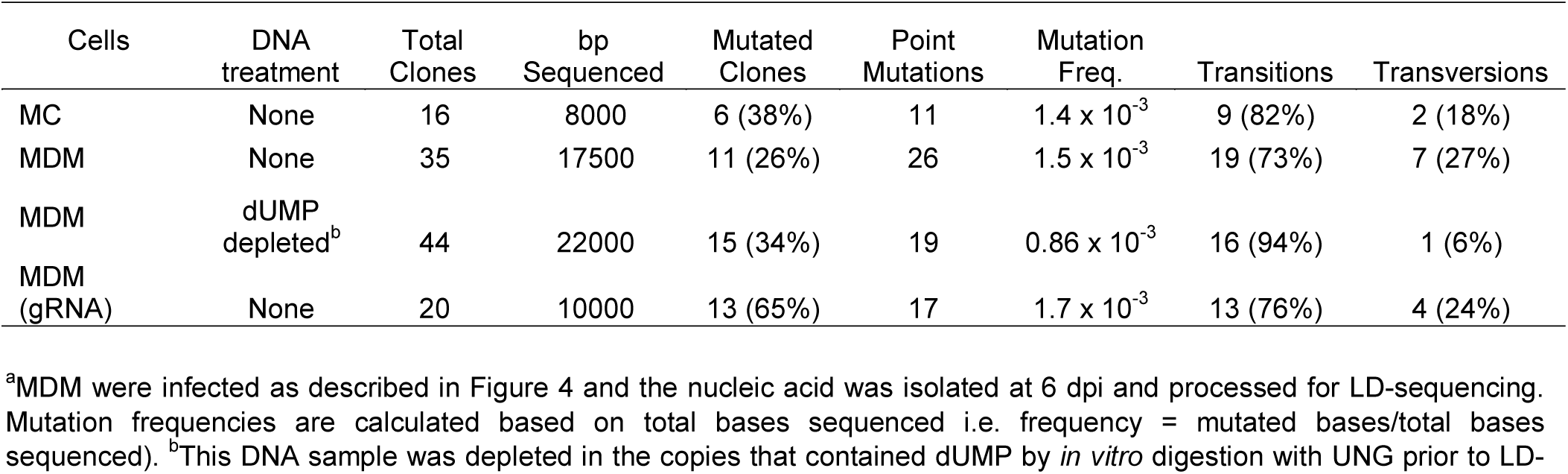
Mutational analysis of HIV proviral DNA and extra cellular viral RNA extracted from infected MC and MDM^a^

| Cells | DNA treatment | Total Clones | bp Sequenced | Mutated Clones | Point Mutations | Mutation Freq. | Transitions | Transversions |
| --- | --- | --- | --- | --- | --- | --- | --- | --- |
| MC | None | 16 | 8000 | 6 (38%) | 11 | $1.4 \times 10^{-3}$ | 9 (82%) | 2 (18%) |
| MDM | None | 35 | 17500 | 11 (26%) | 26 | $1.5 \times 10^{-3}$ | 19 (73%) | 7 (27%) |
| MDM | dUMP depleted<sup>b</sup> | 44 | 22000 | 15 (34%) | 19 | $0.86 \times 10^{-3}$ | 16 (94%) | 1 (6%) |
| MDM (gRNA) | None | 20 | 10000 | 13 (65%) | 17 | $1.7 \times 10^{-3}$ | 13 (76%) | 4 (24%) |

<sup>a</sup>MDM were infected as described in Figure 4 and the nucleic acid was isolated at 6 dpi and processed for LD-sequencing. Mutation frequencies are calculated based on total bases sequenced i.e. frequency = mutated bases/total bases sequenced). <sup>b</sup>This DNA sample was depleted in the copies that contained dUMP by *in vitro* digestion with UNG prior to LD-

**Fig 6.**
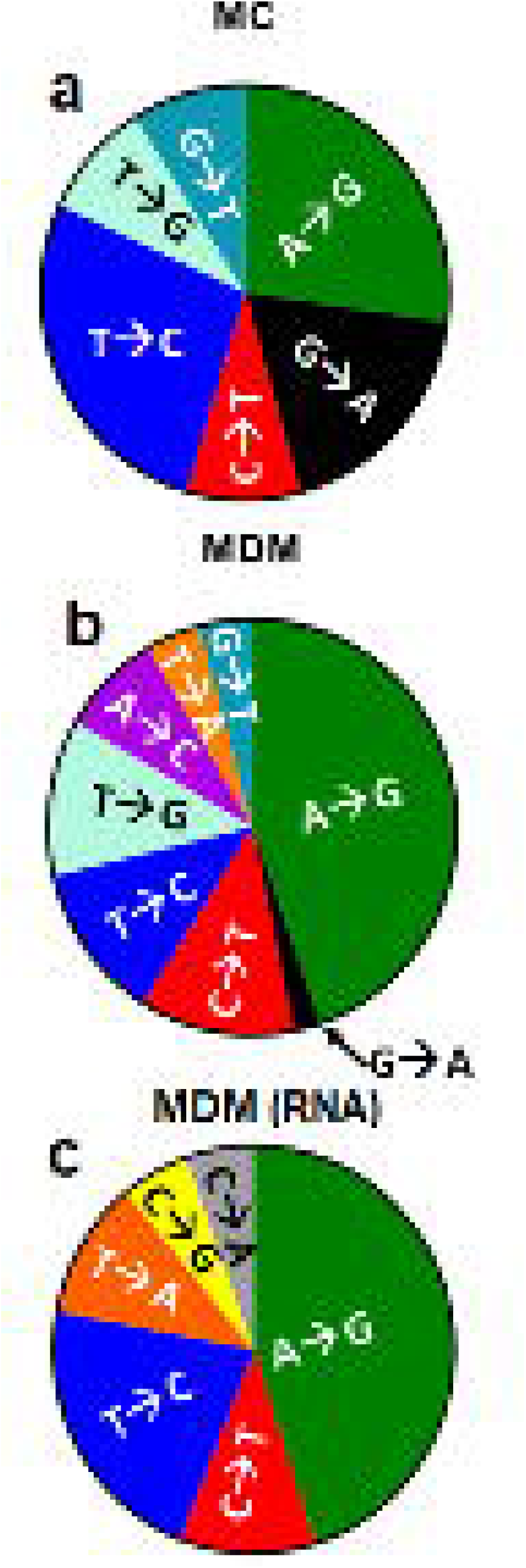
Mutagenic effects of uracil incorporation on HIV proviral DNA and extracellular viral RNA in MDM and MC. Sequences of single proviral clones and extracellular viral RNA from infected MDM and MC. MC and MDMs were infected with equivalent amount of virus (1 pg p24/cell). Seven days after infection total cellular DNA and extracellular viral RNA were extracted from cells and the culture supernatants, respectively. Single viral copies were amplified by limiting-dilution nested PCR and sequenced by the Sanger method. Sequences were aligned to our laboratory reference HIV-1^BaL^ sequence. (**a**) Mutational spectrum of HIV proviral DNA sequences from infected MC. (**b**) Mutational spectrum of HIV proviral DNA sequences from infected MDM (**c**) Mutational spectrum of extracellular viral RNA extracted from infected MDM.

Since the mutational spectrum for integrated virus was indistinguishable for MC and MDM, we chose to selectively sequence extracellular viral RNA produced from HIV-1^BaL^-infected MDM (**Fig 6c**). The viral RNA sequences showed a slightly elevated mutation frequency compared the proviral DNA (1.7 ×10^−3^) and similar percentages of transition and transversion mutations. In addition, about 65% of the viral RNA mutations led to codon changes (**S5 Table**). We recently reported a mutational frequency of 0.6 × 10^−3^ for RNA pol II transcription using linear uracilated DNA templates that were transfected into human UBER deficient Hap1 cells [9]. In this study, the 2.3-fold higher mutation frequency arising from reverse transcription obscures any additional viral RNA mutations that might arise from RNA pol II transcription.

### Selective sequencing of dUMP-depleted DNA fraction

We were interested in whether a different mutational spectrum might result if the fraction of proviral DNA that contained dUMP was subtracted from the total population of proviral DNA before single-molecule amplification and sequencing. To address this question, we performed UNG digestion on the total DNA extracted from infected MDM to remove all of the dUMP-containing copies prior to PCR amplification and then repeated the limiting dilution steps and clonal sequencing (**Fig 7a)**. Although the dUMP-depleted DNA showed a modest 2-fold reduction in the mutation frequency (0.8 × 10^−3^) (**Table 1, S4 Table**), this reduction was not highly significant (*p*-value = 0.16). Despite the insignificant effects on the mutation frequency, the mutation spectrum for the dUMP-subtracted DNA fraction consisted almost entirely of transition mutations, with only a single transversion mutation after sequencing 22,000 bases (**Fig 7a**). We calculate that there is only a 1% probability that a single transversion mutation would have been randomly observed in the dUMP-depleted sample based on Poisson statistics and the measured transversion frequency in the dUMP-rich DNA sample. Thus, transversion mutations appear to be correlated with the DNA fraction that contains dUMP.

**Fig 7.**
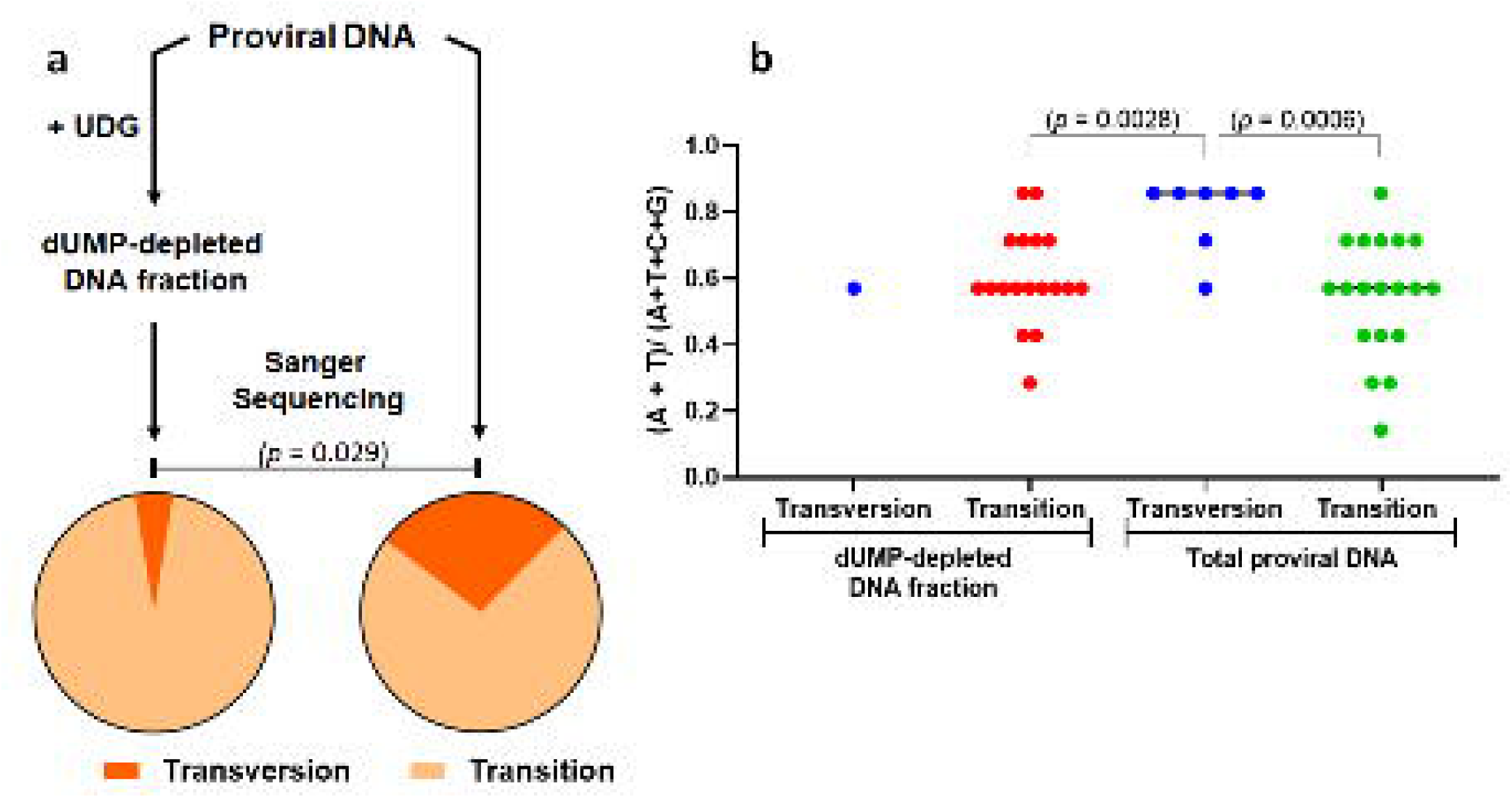
Viral cDNA containing dUMP contains unique (+) strand transversion mutations that correlate with A/T(U) content surrounding the transversion site. MDMs were infected (1 pg p24/cell). Seven days after infection total cellular DNA was extracted from cells. Single copy sequences were determined by limiting-dilution nested PCR and Sanger sequencing. Sequences were aligned to our laboratory reference HIV^BaL^ sequence. (**a**) A population of viral cDNAs was sequenced before and after the removal of dUMP-containing sequences as indicated. Removal of dUMP-containing viral sequences was accomplished by treatment of the total DNA population with UNG prior to PCR amplification and clonal sequencing. HIV proviral DNA sequences where the dUMP fraction was subtracted contained almost exclusively transition mutations (left pie chart), while the total proviral DNA without subtraction showed the same transition mutations but also unique transversion mutations (right pie chart). (**b**) Transversion mutations are associated with loci that contain higher average A/T(U) content. In this analysis, the average A + T(U) content over a window of three flanking bases on both sides of each transversion or transition mutation was calculated. The loci with transversion mutations had a higher average content than the 29 loci that showed transition mutations. *p*-values from an unpaired Student’s *t*-test indicate the statistical significance of the average A + T(U) content in the six base window surrounding the transition and transversion sites.

To explore why DNA clones containing dUMP showed a higher frequency of transversion mutations, we examined the average [A + T(U)] content for 7 mer sequences centered on these transversion sites to explore whether the mutations might be correlated with the density of dUMP incorporation on either the (+) or (-) strand. For comparison, we determined the average [A + T(U)] content of the 7 mer sequences surrounding the transition mutation sites in the dUMP-depleted and total DNA samples (**S2 and S4 Tables**). This comparison showed a statistically higher average frequency of [A + T(U)] near the transversion sites (µ = 0.80) as compared to the transition mutation sites (dUMP-depleted DNA, µ = 0.58; total DNA, µ = 0.55) (**Fig 7b**). The differences in the mean values are significant with *p* values < 0.003. Possible mechanistic implications of this result are discussed below.

### Relative infectivity of HIV-1^BaL^ produced from MC, MDM^EI^ and MDM^LI^

We determined the relative efficacy by which MC, MDM and activated T cells produce virus particles and extracellular vRNA by normalizing the viral output by the average number of proviruses present in each infected cell type (**Fig 8a, b**). Using p24 or extracellular viral RNA levels as the measure for output, there were only modest differences for the three myeloid cell infections at 7 dpi, although MDM that were infected after complete differentiation (MDM^LI^) showed a 2-fold greater yield than MC or MDM^EI^. For comparison, activated T cells infected under identical conditions showed a 5-fold higher viral output than MC (**Fig 8a, 8b**), indicating an intrinsically higher efficiency than any of the infected myeloid cells.

**Figure 8.**
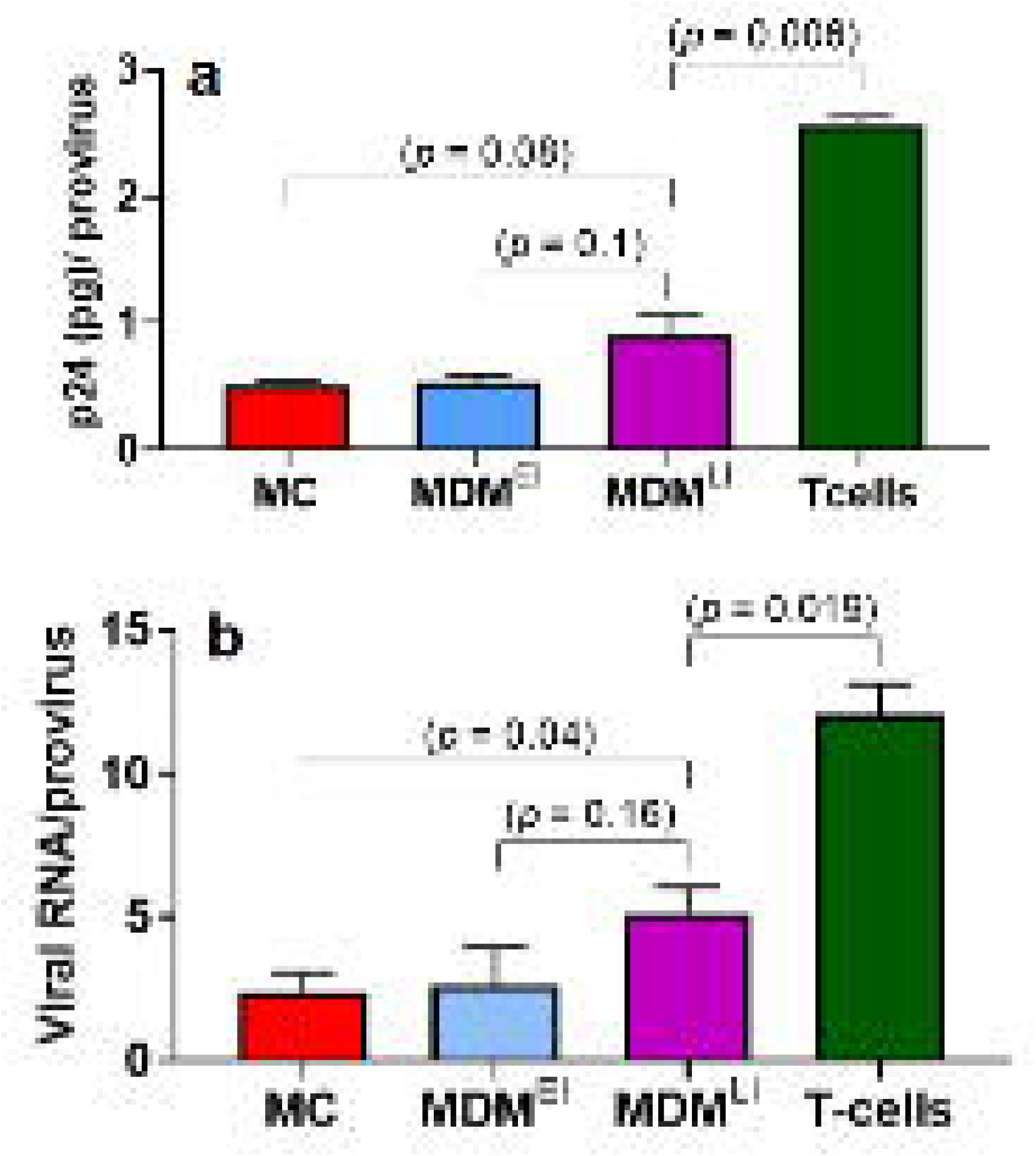
Efficiency of viral particle production from HIV proviruses in MC, MDM and activated T cells. All cell types were infected with Bal virus at moi of 5 (1 pg p24/cell) and seven days after infection culture supernatants and cellular DNA were processed. MC were infected immediately after isolation and then cultured for seven days without differentiation using nonadherent conditions in the absence of M-CSF. MDM^EI^ were infected as freshly isolated MC and then differentiated into MDM under adherent conditions in the presence of M-CSF. MDM^LI^ were infected after differentiation for seven days in the presence of M-CSF. T cell controls were infected and cultured in IL-2 medium for seven days. T20 entry inhibitor (10 µM) was used to prevent reinfection. (**a**) Viral protein 24 (p24) levels in culture supernatants were measured by ELISA and normalized to total proviral copies determined by alu-gag qPCR. (**b**) Extracellular viral RNA was measured using RT-PCR and normalized to total proviral copies determined by alu-gag qPCR. *p*-values from an unpaired Student’s *t*-test are shown.

Finally, we investigated the relative infectivity of HIV-1^BaL^ produced from infected MC, MDM^EI^, MDM^LI^ and activated T cells by collecting culture supernatants at 7 dpi and using equivalent copies of virus to infect MOLT-4/CCR5 target cells (**Fig 9a**). Seven days after infection of the target cells, total cellular DNA was extracted, and LRT copies were quantified using qPCR (**Fig 9b**) and p24 levels were measured in the culture media (**Fig 9c**). Although equal p24 units were used in each of the infections, the target cell LRT copies were about 4-fold greater for infections initiated with viral supernatants obtained from MDM^LI^ as compared to MC, while MDM^EI^ showed an intermediate level of LRT copies. In addition, supernatant p24 levels were 6-fold greater for infections initiated with MDM^LI^. These differences indicate that the fitness of viral particles produced from MC is lower than MDM despite the similar efficiency of production from integrated viruses (**Fig 8**). If we define an infection cycle that begins with viral infection of a MC or MDM and ends with the infection of a new target cell, we calculate an overall 50-fold lower efficiency for MC as compared to a fully differentiated MDM. This comparison between the two cell types is based on the product of the relative efficiencies for virus integration (**Fig 4**), viral particle production from integrated virus (**Fig 8**) and the productive infection of new target cells (**Fig 9**). Using the same calculation, MC are 300-fold less efficient at producing infective virus as compared to activated T cells. It is not clear at this time how much viral dUMP levels or UBER contributes to the lower infection efficiency in myeloid cells.

**Fig 9.**
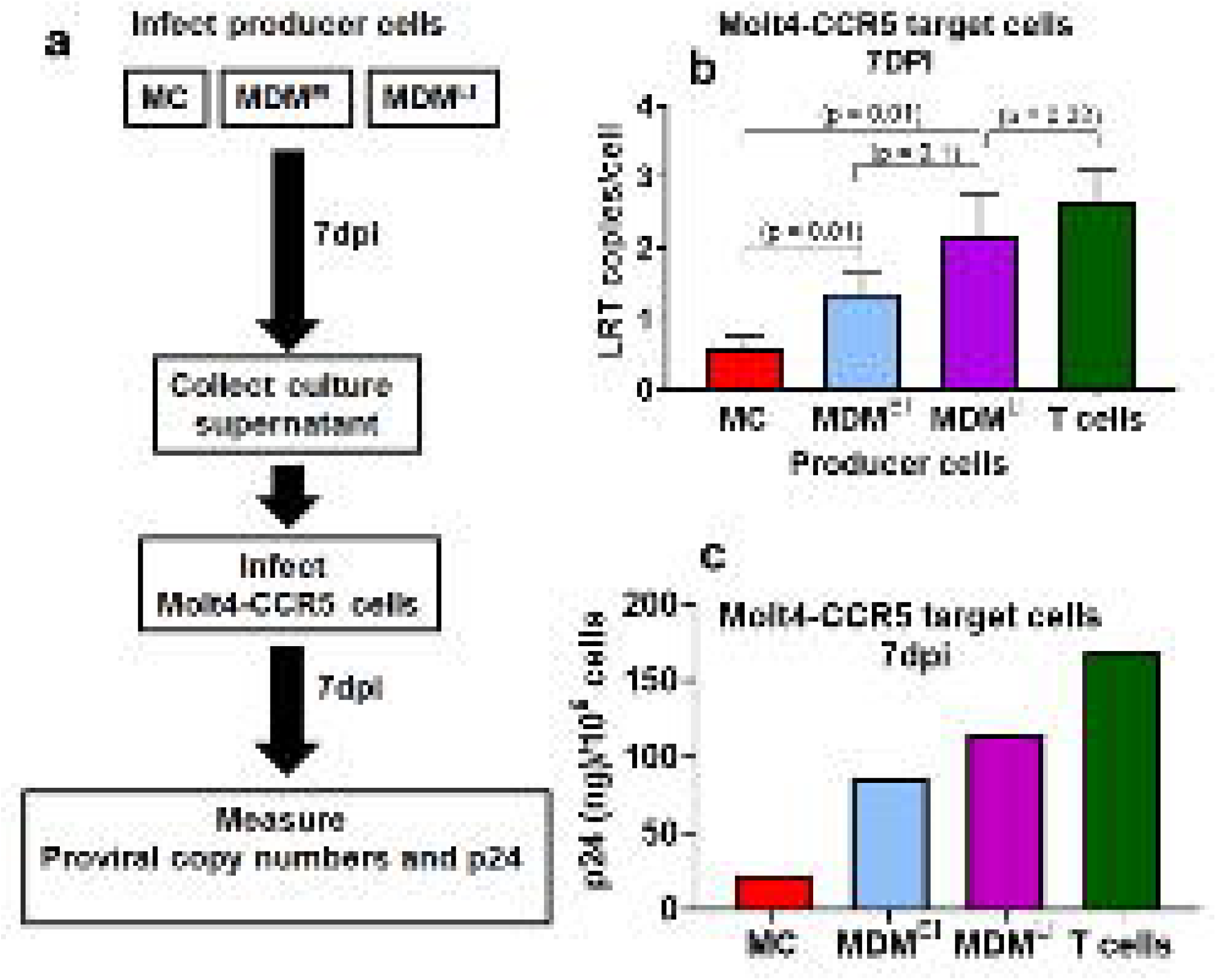
Efficiency of viral particle production from HIV proviruses in MC, MDM and activated T cells. (**a**) Experimental scheme for determining infectivity of virus generated from MC, MDM^EI^ and MDM^LI^ producer cells. (**b**) Viral supernatants from MC, MDM^EI^ and MDM^LI^ and T cell producer cell cultures were collected at 7 dpi and MOLT-4/CCR5 target cells were infected using a 0.1 pg p24/target cell. Seven days after infection of target cells, cellular DNA was extracted and levels of proviral DNA were determined by qPCR. Relative infectivity is shown as LRT copies per target cell. (**c**) p24 levels as measured by ELISA (single donor). *p*-values from an unpaired Student’s *t*-test are shown.

### Expression of exogenous hUNG2 before HIV infection depletes uracilated HIV DNA products

Because of the vanishingly low levels of UBER pathway enzymes detected in MDMs, and especially the very low hUNG2 activity observed in both monocytes and MDMs, we hypothesized that low hUNG2 levels might limit the restrictive effects of the UBER pathway. To test this hypothesis, we over expressed full length human UNG (hUNG2) in MDMs using a doxycycline inducible lentiviral transduction system. The inducible system allowed us to test whether hUNG2 expression had a greater effect before HIV infection, or alternatively, after HIV had integrated into the MDM genomic DNA. Control experiments demonstrated the presence of high hUNG2 activity in cell extracts prepared from transduced MDM at days 1 and 3 post induction, but no activity in the absence of doxycycline induction (**S5 Fig**).

We first tested the effect of inducing hUNG2 expression prior to infection with VSVG pseudo-typed HIV^pNL4-3^ virus particles capable of only a single round infection. In this experiment, fully differentiated MDM were first transduced with the hUNG2 expressing lentivirus at an MOI of 5 (0.1 pg p24 antigen/cell) and expression of hUNG was induced using 1μg/ml of doxycycline three days after transduction, followed by infection with HIV^NL4-3^ one day later (MOI = 0.5) (**Fig 10a**). We followed the HIV provirus copy number and the fraction of total provirus that contained dUMP using alu-gag Ex-qPCR. These measured outcomes were compared with those of an uninduced control infection (blue bars, **Fig 10a**), as well as an infection with HIV^pNL4-3^ in the absence of any prior lentiviral transduction (green bars, **Fig 10a**). With pre-infection induction of hUNG2 expression (red bars, **Fig 10a**), there was a 50% decrease in the provirus copy number between 1- and 7-days post HIV^pNL4-3^ infection and the fraction of proviruses containing dUMP was at the limit of detection for the Ex-qPCR method (< 0.2). In contrast, the no induction and HIV^NL4-3^ only control infections showed a stable or slightly increasing proviral copy number over the same time period and the fraction of proviruses containing dUMP increased from ∼0.4 to 0.8. Similar results were obtained using LRT primers for Ex-qPCR (**S6 Fig**). As discussed above and previously [4,27], the reduction in proviruses as a result of UNG overexpression is never complete because the MDM exist as a mixed G1-G0 population where only about 40-50% of the total proviruses contain dUMP. In contrast, the essentially complete absence of viral dUMP at one to seven days post infection as measured by alu-gag Ex-qPCR clearly indicates cellular hUNG2 excised the uracils prior to viral integration. These results indicate that when hUNG2 is abundant in the target cell prior to HIV infection, uracilated viruses can be efficiently destroyed before integration.

**Figure 10.**
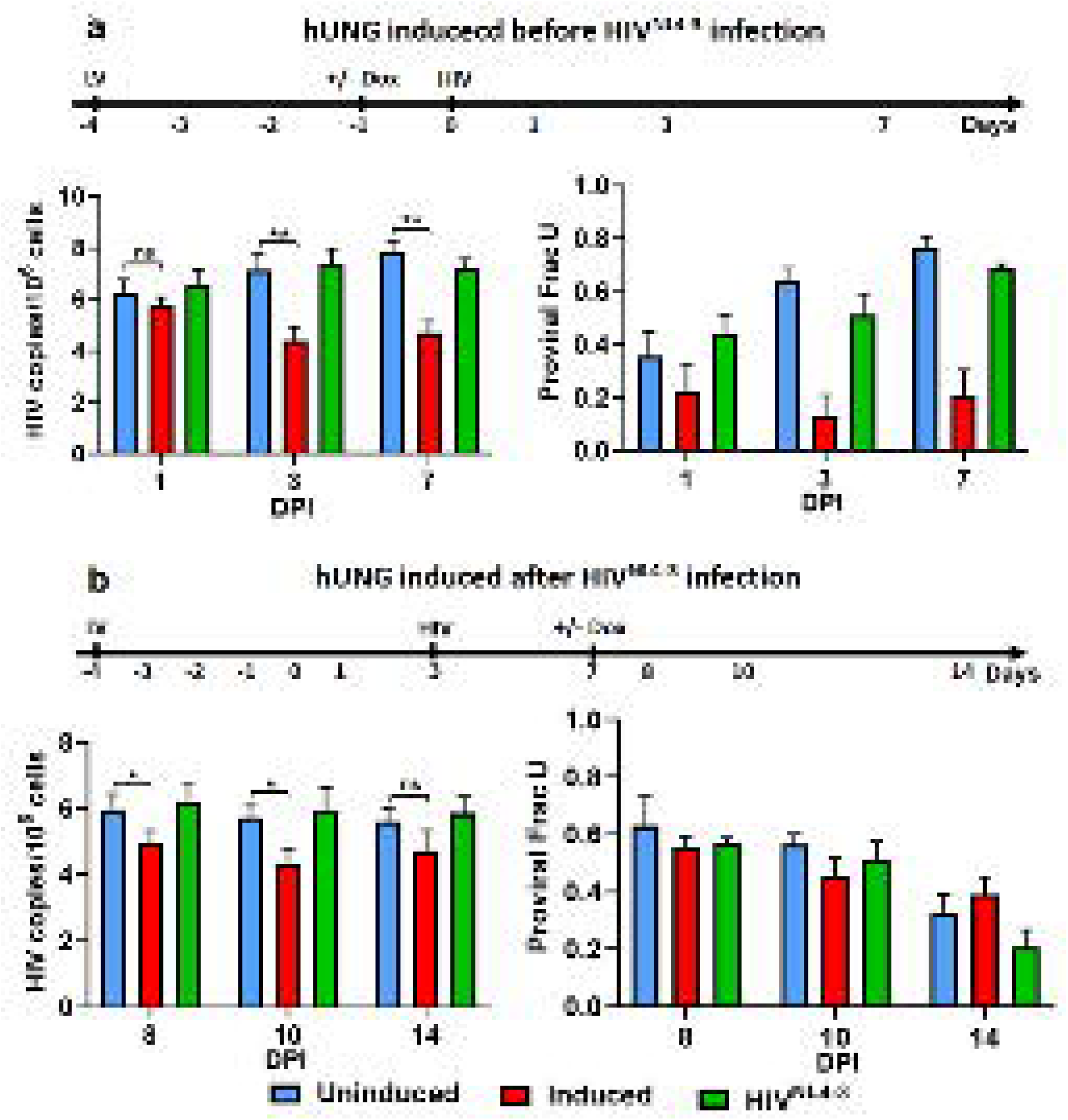
Exogenous expression of hUNG2 depletes uracilated HIV DNA products. Exogenous full-length hUNG2 was over expressed in MDM under control of a doxycycline (Dox) inducible promoter by lentiviral transduction (moi = 5). The timelines above each panel indicate the experimental course and the points where hUNG2 expression was induced relative to infection with HIV^NL4-3^. (**a**) hUNG2 expression was induced with doxycycline (1 µg/ml) one day before HIV^NL4-3^ infection. Proviral copy numbers (left panel) and the fraction of HIV proviral DNA that contained dUMP were measured at indicated times (FracU, right panel). (**b**) Expression of hUNG2 was induced 7-days after infection with HIV^NL4-3^ in order to allow HIV to fully integrate. Proviral copy numbers (left panel) and the fraction of HIV proviral DNA that contained dUMP were measured at indicated times (FracU, right panel). Uninduced (blue) and no lentiviral infection (green) controls were also performed.

In a second experimental protocol, hUNG2 induction was delayed until 7 days after HIV^NL4-3^ infection to allow most of the HIV cDNA to integrate (**Fig 10b**). Under this infection scenario we saw a smaller decrease in HIV proviral DNA between 8- and 14-days post-infection (one to seven days post-induction) as compared to the pre-infection induction of hUNG2 shown in **Fig 10a**. In addition, a higher fraction of proviruses contained dUMP and these proviruses slowly disappeared over one to seven days post-induction. In fact, the rate of decrease of dUMP-containing proviruses was not significantly different for the hUNG2-induced condition and the uninduced controls (**Fig 10b**). This indicates that the rate-limiting step for post-integration excision of uracils does not involve hUNG2 and may instead involve remodeling of chromatin.

## DISCUSSION

dUTP-mediated innate immunity was first suggested for β-retroviruses, non-primate lentiviruses, and endogenous retroviruses [30–33]. These viruses have all captured a host dUTPase gene during viral evolution, which is a powerful enzyme that degrades dUTP to dUMP and PP_i_, promoting infection of macrophages by these viruses. Although HIV-1 does not encode for dUTPase, it can infect human monocytes and macrophages even though the ratio dUTP/TTP is large. One strategy that HIV uses to replicate in this environment is to code for the accessory protein vpr, which is used to target hUNG2 for ubiquitination and proteasomal degradation [16,34], suggesting that incorporation of dUTP is tolerated but that the antiviral function of hUNG2-catalyzed uracil excision must be overcome. Consistent with this phylogenetic comparison of different retroviruses, our previous work established that this dUTP-mediated restriction pathway works most potently when target cell dUTPase expression is low, dUTP is abundant, and nuclear hUNG2 is highly expressed so as to efficiently degrade viral cDNA [3,4]. When any of these elements are absent or diminished, the effectiveness of the restriction pathway is diminished, which likely explains diverse reports in the literature concerning the impact of this pathway [31,35–37]. Accordingly, quantification of cellular [dUTP]/[dTTP] and UBER enzyme expression levels are key to understanding the impact of this pathway in MC and MDM.

### Implications of UBER activity and [dUTP]/[dTTP] in MC and MDM

Since MC differentiate into macrophages, it is possible that a macrophage originally infected at the MC stage of differentiation might produce different amounts and sequences of virus compared to a macrophage that was infected after complete differentiation (**Fig 1**). This question pertaining to the order of infection is relevant because MC containing uracilated viral DNA have been detected in blood samples of HIV patients on ART who are in full remission (10 – 600 infected MC/10^6^ cells)[4,19]. This observation is intriguing because peripheral blood MC have a limited half-life of about 2 days in circulation and in this limited period infection must occur. After infection, MC either migrate to lymphoid tissues and differentiate into macrophages or are eliminated [20]. We speculate that under virally repressed conditions MC are most likely infected while passing through a tissue reservoir containing HIV infected cells and then enter back into circulation via retrograde migration before dissemination to other tissues [22,23].

Here we have demonstrated the kinetic competence of the MC infection pathway *in vitro* through the detection of viral cDNA products in MC within a relevant time frame (**Fig 4c-e**). Since MC have high dUTP levels, but the entire UBER pathway is almost absent with the exception of APE1 (**Fig 3**), dUMP-containing proviruses likely persist throughout the MC lifetime. Thus, infected MC harboring uracilated viral DNA may successfully migrate to a tissue compartment and differentiate into macrophages. Once differentiated into macrophages, UBER activity is upregulated and dUMP can be slowly excised and at least partially replaced with dTMP. This mechanism is supported by the observed slow disappearance of viral dUMP over 28 days in MDM, with no change in proviral copy number, suggesting that dUMP is being replaced by dTMP via UBER (**Fig 4b**). This mechanism requires that the dTTP concentration is competitive with dUTP to prevent futile cycling (futile cycling is the excision of dUMP and its reincorporation by DNA pol). The measured [dUTP]/[dTTP] ∼1 in macrophages in our *in vitro* culture conditions is compatible with a repair mechanism that allows slow nucleotide replacement over time (**Fig 2**). In contrast, the [dUTP]/[dTTP] in the range 20 to 60 as found in two previous studies would effectively prevent repair, and excision of such densely spaced uracils by hUNG2 would likely lead to fragmentation of the viral DNA as previously indicated [3,4,24]. If the [dUTP]/[dTTP] ratio is variable *in vivo*, it will have an impact on the efficiency of HIV infection of myeloid lineage cells, with outcomes ranging from slow repair to viral destruction depending on the dUMP density and hUNG2 expression levels.

### UBER may occur after switching of macrophages from a G_0_ to G_1_ cell cycle state

Mlcochova *et al* reported that non-dividing MDM exist in two interconvertible cell cycle populations, a minor population that exhibits markers of G_1_ and is DNA repair competent and a predominant population that is G_0_-like and deficient in the expression of many DNA repair proteins and DNA replication components [27]. The two states can also be detected by infection with an HIV construct containing an eGFP expression cassette [4,27,38]. In this case, the restrictive G_0_ population shows little or no eGFP fluorescence, even though viral cDNA is present in nearly every cell, while the permissive G1 population shows high eGFP expression [4,38]. These observations of Mlcochova *et al* parallel our previous measurements using a fluorescent reporter virus [4]. In our previous study we sorted infected MDM into GFP positive and negative populations and found that the GFP positive (G_1_) population did not contain any detectable dUTP and the viral DNA was free of dUMP (4). In contrast, the GFP-negative (G_0_) population contained high [dUTP] and the viral DNA also contained high levels of dUMP [4]. More recent work on the effects of dUMP on transcription of DNA templates by RNA pol II establishes that the differences in GFP expression between the two populations is at least partly due to transcriptional repression by dUMP [9]. Thus, these combined findings suggest that UBER may occur after an infected G_0_ macrophage transiently moves to the G_1_ state where the nucleotide pool composition and DNA repair capacity can facilitate dUMP removal and replacement. We speculate that small molecules that target the G_0_-G_1_ checkpoint might be useful to induce the transition to the G_1_ state and potentially lead to inactivation of heavily uracilated proviruses by upregulation of UBER.

The realization that bulk MDM consist of two cell populations with respect to dUTP levels and UBER capacity further informs our understanding of the observation that only a fraction of total viral cDNA molecules contain dUMP. For instance, assuming a homogenous MDM population, it is difficult to rationalize why only ∼60% of total proviruses contain dUMP within a 1200 base pair amplicon (**Fig 4b**). This fraction is enigmatic because with [dUTP]/[dTTP] ∼ 1 and the inability of reverse transcriptase to discriminate between dTTP and dUTP (**Fig 2**), every 10 bp of viral DNA should contain dUMP on both strands and all proviruses should contain dUMP. Thus, the low fraction of uracilated proviruses in MDM is consistent with the presence of two cell sub-populations and only one of the populations (G_0_) contains high [dUTP]/[TTP]. In contrast, the observation that nearly all copies of HIV contain persistent dUMP during infection of MC suggests that the G_1_ population does not exist to a significant extent before differentiation into MDM. Consistent with this model, MC show virtually no eGFP expression upon infection with a GFP reporter virus even though proviral DNA is present in nearly every cell (**S4 Fig**).

### Potential mechanisms for mutational effects of dUTP incorporation by reverse transcriptase

There are surprisingly few studies of the mutational events during HIV infection of MDM, and to our knowledge, no previous reports of HIV sequences derived from infection of MC. The most complete and direct comparison of HIV mutations arising during single round infection of T cells and MDM is the next-generation sequencing study of full-length HIV genomes by Cromer et al [39]. In this study, the reported overall mutation frequency was ∼1 × 10^−4^ for both cell types. Their reported HIV mutation spectrum for MDM infections (G⍰A = 35%, A⍰G = 18%, A⍰T = 17%, A⍰C = 11%) does not differ significantly from the profile obtained in our selective sequencing of a 500 bp amplicon covering the V3 and V4 regions of *env*, which is known to be highly variable (**Fig 6b**)[40,41]. However, our mutation frequency for this select region (1.4 × 10^−3^, **Table 1**) is significantly higher than the average value previously reported by Cromer et al for the entire HIV sequence. This result is not unanticipated because we have previously reported that the mutation frequency for HIV DNA and RNA isolated from infected MDM is variable across the HIV genome [4]. In particular, the LTR region has very low levels of mutations [4].

One significant observation in our sequencing studies was that transversion mutations in proviral DNA were not detected in DNA samples where the dUMP fraction was subtracted by UNG digestion (**Table 1, Fig 6c**). This result indicates that the mutational profile for misincorporation of nucleotides by reverse transcriptase is indirectly impacted by the presence of dUMP in the (-) or (+) strand sequences. Although the detailed mechanistic basis for this outcome is not known, the sites of these transversion mutations are especially AT(U) rich (**Fig 7b**), suggesting that uracils on either strand may be involved (**S2 Table**). There is currently no information about the stability or dynamics of dUMP base pairs in the context of RNA/DNA hybrids, the possible sequence contexts that might give rise to mutational hotspots, or the activity of reverse transcriptase under such conditions. Relevant to the current findings, previous work on dUMP/A base pairs in duplex DNA has shown increased base pair dynamics and decreased thermodynamic stability [28,29,42]. The increased dynamics has been studied using NMR imino proton exchange and attributed to the reduced electron density of the base, which leads to poorer stacking interactions with neighboring bases [29]. Such effects in the context of RNA/DNA hybrids containing densely spaced dUMP substitutions are anticipated but require detailed biochemical studies to confirm. In this regard, we recently reported reduced fidelity of T7 RNA polymerase during transcription of RNA using a DNA template that contained dUMP [9]. These mutations were also explained by indirect effects of dUMP arising from destabilization of the template-primer, which increased strand slippage and realignment errors during polymerization. Similar observations with reverse transcription now suggest this proposed effect of dUMP may be general.

A further aspect of interpreting the mutational spectrum that arises during dUMP incorporation is whether the errors occur during minus strand or plus strand synthesis. Two observations that favor a predominantly minus strand error mechanism are (i) the known lower fidelity of reverse transcriptase when using an RNA template [43], and (ii) the complete absence of base mismatches detected during DNA sequencing (for instance, an original G/U base mismatch should yield Sanger strand reads consisting of a mixture of G/C and A/T at that site after PCR amplification). The occurrence of predominantly minus strand errors is suggested because a mismatch generated during (-) strand synthesis would always be removed during degradation of the plus strand RNA by HIV RNaseH, whereas mismatches occurring during plus strand synthesis would remain. Although DNA repair could repair plus strand mismatches, the low repair capacity of nondividing cells makes this unlikely compared to the simple minus strand mechanism. Consistent with this idea, a recent deep sequencing study concluded that minus strand deamination events catalyzed by A3G were only inefficiently repaired by UBER or mismatch repair in T cells, which have much greater repair activity than MDM [44].

## CONCLUSIONS

These in vitro results inform on the previous detection of dUMP in both alveolar macrophages and peripheral blood monocytes of HIV patients on ART [4]. We find that the kinetics for MC infection is compatible with their lifetime *in vivo* and their near absence of UBER activity is consistent with the retention of viral dUMP at high levels at least until differentiation into macrophages, where UBER becomes possible. Thus, the fate of uracilated proviruses would appear to center on the expression levels of UBER enzymes in different cellular environments and whether infected macrophages can enter the repair competent G_1_ state. A full understanding of these complexities in the context of *in vivo* derived macrophages is therefore essential. Our current understanding suggests that potential therapeutic opportunities may exist for eradicating densely uracilated proviruses if UBER could be activated under conditions where subsequent repair is interrupted pharmacologically.

## Supporting information

Supplemental References

## Acknowledgments

The authors acknowledge the support of the Johns Hopkins Institute for Clinical and Translational Research team for recruiting, consenting and performing blood draws from healthy subjects.

## Supporting information

**S1 Supplemental Methods.** RT-PCR measurements of T cell receptor RNA, single nucleotide polymerase extension assay, western Blotting, measurement of mRNA expression levels of UBER enzymes using RT-qPCR, uracil content of viral DNA, RT-qPCR of extracellular viral RNA, activity of HIV reverse transcriptase with dUTP and dTTP substrates, sequencing of single viral reverse transcripts and genomic RNA copies, over-expression of hUNG in MDMs, gel-based hUNG activity assay

**S1 Fig. Measurement of dTTP and dUTP levels in the Hap1 dividing cell line.** The single nucleotide extension assay was used to establish the differences in dUTP/dTTP between MDM, MC and HAP1 dividing cells. The procedure is described in Methods. The gels show extension reactions in the presence and absence of dUTPase. The indicated fold serial dilutions of the dNTP extracts establish that extension is within the linear range of our assay (0.05-0.9 fraction substrate extended). The total [dTTP + dUTP] pool was 68 ± 6 pmol/million cells which was comprised entirely of dTTP (64 ± 4 pmol/million cells).

**S2 Fig. Determination of UBER mRNA expression levels in MDMs and comparison with the HAP1 dividing cell line. (a)** Relative expression of UBER mRNA of MDMs (blue) and HAP1 (red) cells. Relative expression is with respect to 18s ribosomal RNA. **(b)** Fold differences in UBER mRNA expression levels in MDM relative to HAP1 dividing cells. Total RNA was extracted from MDMs after seven days differentiation from MC. All qPCR reactions were done in triplicate.

**S3 Fig. Morphological and granulation differences between MDMs and MCs. (a)** Flow cytometry analysis of monocytes immediately after purification. (**b**) Flow cytometry analysis of monocytes cultured in suspension under non-adherent conditions for seven days. (**c**) Flow cytometry analysis of fully differentiated MDM (cultured for 7 days under adherent conditions in the presence of M-CSF). (**d**) Light microscope image of monocytes immediately after purification (20x magnification, 5x zoom). **(e**) Light microscope image of monocytes after 7 days of culture under non-adherent conditions in the absence of M-CSF (20x magnification, 5x zoom). (**f**) Light microscope image MDM after 7 days of culture in the presence of M-CSF (20x magnification, 4x image reduction

**S4 Fig. HIV-eGFP expression in MC and MDM.** (**a**) MC were infected with HIV^NL4-3(eGFP)^ immediately after isolation at an moi of ten. At 7-days post infection, eGFP expression was measured by flow cytometry. Even though GFP fluorescence is very low, viral reverse transcripts are abundant (main text). (**b**) Fully differentiated MDM were infected with HIV^NL4-3(eGFP)^ at an moi of ten. At 7-days post infection, eGFP expression was measured by flow cytometry.

**S5 Fig. Activity of lentiviral transduced hUNG2 in MDM cell extracts under uninduced and induced conditions.** MDM were transduced with a lentiviral construct containing full length hUNG2 (pCW.57.1.FL.UNG) at MOI of 5 (0.1 pg p24/cell). Total cell extracts were prepared at days 1 and 3 post-transduction and protein concentrations were determined by the Bradford assay. hUNG activity in cell extracts was determined by gel-based UNG activity assay using equal amounts of total protein and a 19-mer uracil-containing ssDNA substrate with a FAM label on the 5’-end (see Methods). Excision of uracil results in a 5’-FAM labeled 9mer product band.

**S6 Fig. Effect of over expression of hUNG2 in MDMs on total HIV DNA.** Fully differentiated MDM were first transduced with inducible lentiviral construct expressing full length hUNG at an MOI of 5 (0.1 pg p24/cell) and 3 days later induced with doxycycline (1ug/ml). 1-day after induction, MDM were then infected with HIV^NL4-3^ single round virus at MOI of 0.5 (0.05 pg p24/cell). Total DNA was extracted at days 1, 3 and 7 and (**a**) LRT copies and (**b**) Frac U were measured by Ex-qPCR.

**S1 Table.** Primer and molecular beacon probe sequences for viral DNA sequences and RT-PCR measurements of UBER enzyme mRNA expression (5’→3’)

**S2 Table.** Clonal mutation analysis of HIV proviral DNA isolated from infected MDM at 7 dpi **S3 Table.** Clonal mutation analysis of HIV proviral DNA isolated from infected MC at 7 dpi

**S4 Table.** Clonal mutation analysis of dUMP-depleted HIV proviral DNA isolated from infected MDM at 7 dpi

**S5 Table.** Clonal mutation analysis of extra cellular viral RNA extracted from infected MDM culture supernatants at 7 dpi^a^

