## Supplemental Figure 1 for "Deficient uracil base excision repair leads to persistent dUMP in HIV proviruses during infection of monocytes and macrophages"

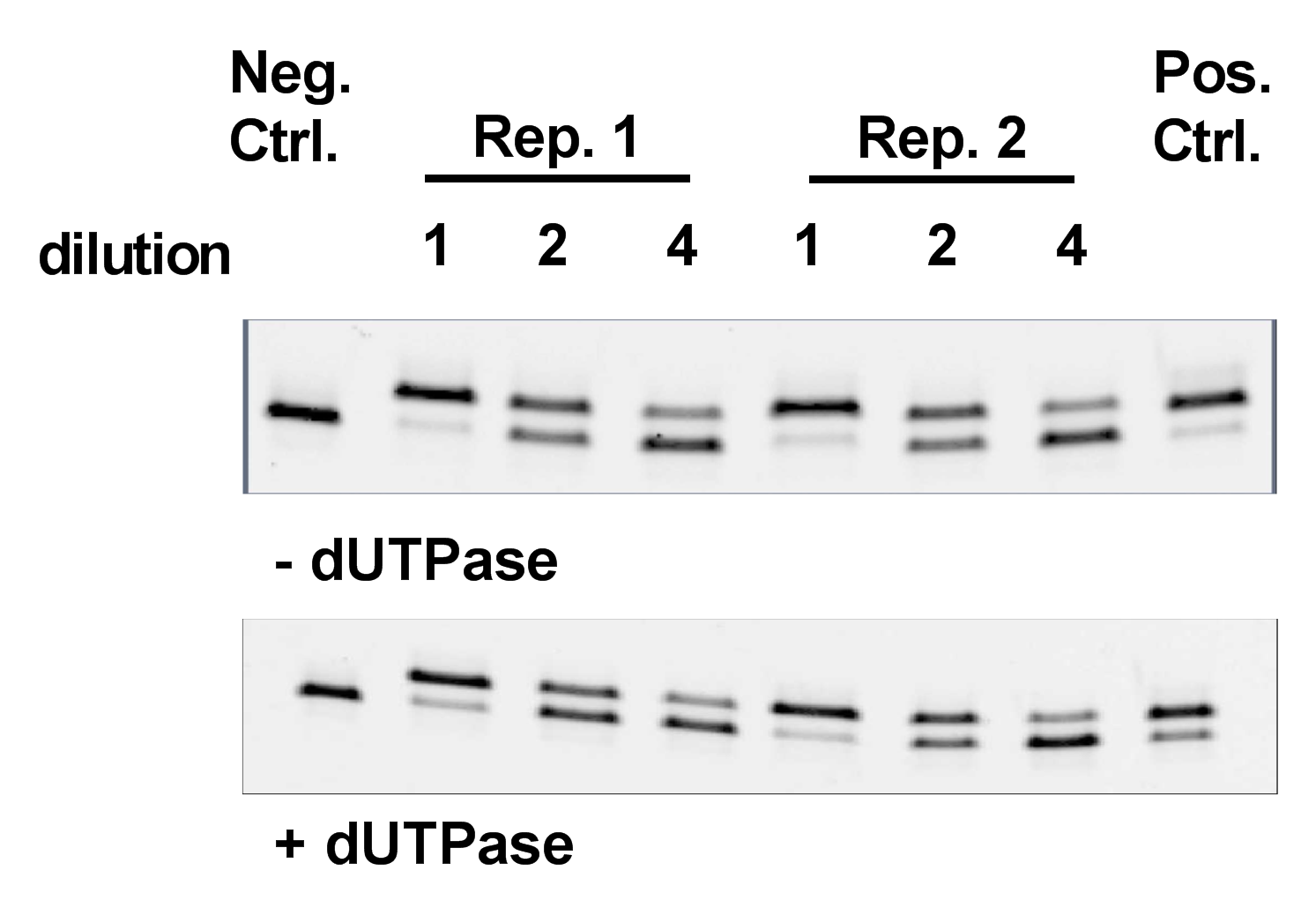

**S1 Fig. Measurement of dTTP and dUTP levels in the Hap1 dividing cell line.** The single nucleotide extension assay was used to establish the differences in dUTP/dTTP between MDM, MC and HAP1 dividing cells. The procedure is described in Methods. The gels show extension reactions in the presence and absence of dUTPase. The indicated fold serial dilutions of the dNTP extracts establish that extension is within the linear range of our assay (0.05-0.9 fraction substrate extended). The total [dTTP + dUTP] pool was 68 ± 6 pmol/million cells which was comprised entirely of dTTP (64 ± 4 pmol/million cells).
