## Supplemental Methods for "Deficient uracil base excision repair leads to persistent dUMP in HIV proviruses during infection of monocytes and macrophages"

**S1 Supplemental Methods**

**RT-PCR measurements of T cell receptor RNA.** Monocytes were purified by negative selection using pan monocyte purification kit (Miltenyi). The remaining cells in the column (PBMC – MC) were also eluted and a million cells were used from each cell pool to extract RNA. One tenth of the total RNA (10^5^ cells) was used to prepare cDNA and quantified using qPCR against a standard curve developed using a plasmid that contains the variable region of the TCR-β. Primers used for both RT and qPCR are indicated in **Table S1**. The contamination from T cells is determined by dividing the TCR copy number measure in the purified monocytes to that in the PBMC - MC cell population (>90% T cells). The expression level of the TCR mRNA in the PBMC – MC fraction was ~10 copies per cell.

**Single nucleotide polymerase extension assay.** The extension reactions (12 or 100 μL) were performed at 37 °C for 40 min and included ~6-1200 fmol of total dNTP, 120 or 1200 fmol of DNA template-primer for extracts obtained from non-dividing or dividing cells, respectively, and 2 units of RTase (one unit converts 1 nmol of labeled dTTP into acid-insoluble material in 20 min at 37°C, pH 8.3). The substrate and the *n* + 1 product of the extension reaction were separated on a 15% denaturing polyacrylamide gel and detection was achieved by following the signal provided by a 5′-fluorescein label located on the primer strand. To remove dUTP from the extracts and specifically measure the dTTP present, the extract was depleted of dUTP by the addition of 10 pmol of dUTPase for 30 min prior to performing the extension reaction. The extension buffer consisted of 20 mM Tris-HCl, pH 8.0, 100 mM KCl, 5 mM MgCl_2_, 2 mM DTT, and 0.1% BSA. Extensive validation experiments were performed using [dUTP + dTTP] mixtures with known dUTP/dTTP 1:1 ratio to establish a detection limit of >6 fmol dNTP and a linear correlation between the known dNTP concentrations and the ratio determined from the SNE assay. No assay interference from addition of up to 1 mM ribonucleotide triphosphates was observed.

**Western Blotting.** Extracts for western blotting were prepared using denaturing conditions. Approximately 4 million MC or 2 million MDM were collected from culture plates and washed three times with PBS before the addition of 100 μL of denaturing lysis buffer to the pellet (CellLytic M buffer + 0.2 % w/v SDS + 10 mM DTT). Protein content in the extracts was measured using the BCA assay (Pierce). The sample concentrations were adjusted to 2 μg/μL using the same denaturing-lysis buffer and frozen at -20 °C until use. Ten μg (5 μL) of each cell extract was loaded on a 4 to 12% NuPAGE Bis-Tris gradient gel (Novex) and electrophoresis was performed using 1x MES-SDS buffer (Novex). Proteins were transferred to a Biorad-select PVDF membrane (Bio-Rad) using 1-step transfer buffer (Pierce). The membranes were stained using SuperSignal West Pico PLUS Chemiluminescent Substrate kit (Thermo**)** for 40 s. Membranes were placed on a transparancy sheet and imaged using a Biorad ChemiDOC XRS system for 15 to 60 s to obtain images in the linear exposure range. The intensity of the individual bands corresponding to a tubulin loading control or the protein of interest were obtained by densitometric analysis in Quantity One Basic. Expression levels were measured by densitometry and normalized to the tubulin reference. We confirmed that the staining was in the linear range of detection for each antibody by increasing the sample load by a factor of two. Primary antibodies for SAMHD1 (ab67820), APE1 (ab137708), pol-β (ab26343), and tubulin (ab15246) were purchased from Abcam. The antibody against DNA ligase 3 (LigIII) was from GenTex (GTX103172) and the anti-human APOBEC3G polyclonal antibody (also detects ABOBEC3A) was obtained from the NIH AIDS Reagent Program. The secondary antibodies for goat anti-rabbit HRP (ab97080**)** and goat anti-mouse HRP (ab97040**)** were obtained from Abcam. The specificities of the antibodies for SAMHD1 have been confirmed in SAMHD1 KO and Hap1 SAMHD1 overexpression cells, the base excision repair enzymes were validated by their strong staining in dividing cells and weak or non-existent staining in non-dividing cells. We used a highly-specific and sensitive hUNG enzyme activity assay (see main text) to determine the low levels of hUNG present in MDM and MC cell extracts, which was further confirmed using RT-PCR quantification of mRNA expression levels.

**Measurement of mRNA expression levels of UBER enzymes using RT-qPCR.** Total genomic RNA was isolated from MDM or HAP1 cells using Trizol Reagent (Life Technologies) following the manufacturers protocol. RNA (~3 μg) was treated with 2 units turbo-DNase (Invitrogen) in 1X turbo-DNase buffer for 30 minutes at 37 C, to remove DNA carryover and then purified using Qiagen RNEasy kit. The RNA concentration was determined using UV absorbance at 260 nm. To generate cDNA library from the isolated total RNA using random primers and High-Capacity cDNA Reverse Transcription Kit (Applied Biosystems) was used with RT thermal program of 25 ^o^C for 10 mins, 37 ^o^C for 2 hours and 85 ^o^C for 5 mins. Quantitative real-time PCR with SYBR green detection was used to calculate the comparative concentration of each gene of interest (UNG, APE1, LigIII, Polβ, SAMHD1, A3A, and A3G) using comparative quantification analysis built in the Rotor-Gene Q Series Software (**Fig. S2**). 18S ribosomal RNA was used as the calibration standard. Thermal cycling conditions for qPCR consisted of 95 C for 5 min, and 40 cycles of denaturation at 95 C for 10 sec and annealing and extension at 60 C for 30 sec. All primers are listed in **Table S1.**

**Uracil content of viral DNA.** Uracil content of viral DNA was determined either using excision droplet digital PCR (Ex-ddPCR) or a similar qPCR method (Ex-qPCR) as previously described (1) with some modifications. For Ex-ddPCR, genomic DNA was first fragmented for 1 h at 60 °C using the endonuclease BSAJ-1 (1 U) in Cutsmart buffer (NEB). To determine the uracil content of the viral DNA, one-half of the sample (~50 ng) was digested with 0.125 units of UDG (New England BioLabs, M0280S) for 30 mins at 37 ^0^C, rendering any uracil containing amplicons inert to PCR amplification. DNA remaining in the UDG-treated or mock-digested DNA samples was quantified using alu-gag nested ddPCR. As described above, the first PCR amplification was performed post-UDG digestion using *alu* forward and *gag* reverse primers. This first PCR product was diluted 20-fold and 5 μl of the diluted PCR product was used as template for droplet amplification using the ERT forward and reverse primers. Each 20 μL ddPCR reaction contained 10 μL Probe Supermix (Bio-Rad), 900 nM each primer, 250 nM ERT probe, and template DNA from the first PCR reaction. PCR mix was loaded into the Bio-Rad QX100 droplet generator and droplets were generated following the manufacturer’s protocols. The contents were transferred to a 96-well reaction plate and sealed with a pre-heated Eppendorf 96-well heat sealer for 5 seconds. Amplification was performed separately in a thermal cycler with the following thermal program: 95 °C for ten min, followed by 44 cycles of 95 °C for 15 s, 60 °C for one min, followed by a final step of heating to 95 °C for ten minutes. The final high-temperature cycle serves to cure the droplets. For each step, the ramp cycle time was decreased by 0.5°C/s. Amplified samples were transferred to a Quanta droplet reader (Bio-Rad) for analysis. Using a commercial primer-probe mix (Bio-Rad) the copy number of the human RNaseP gene (*RPP30*) was measured in the same reaction and used as a reference standard. Ex-ddPCR data were analyzed using the QuantaSoft program (Bio-Rad) and the fraction of the amplicons that were uracil-free was calculated using [eq 1](https://elifesciences.org/articles/18447#equ1) (1).

| FracU^DNA^ = | [positive droplets (no UDG) − positive droplets (+ UDG)] | (1) |
| --- | --- | --- |
|  | [positive droplets (no UDG)] |  |

In some cases excision-qPCR was used to determine uracil-containing fraction of viral DNA. As with the Ex-ddPCR procedure, the sample is split into two equal portions and one portion is treated with UDG. Briefly, 0.125 units of UDG (NEB) was added into the Qiagen Rotor Gene Probe PCR master mix to excise uracils from viral DNA. The PCR thermocycler reaction was modified to include the UDG reaction time and heat-cleavage of the resulting abasic sites. Thermocycler program we used for this reaction was: 37 ^o^C for 30 min (UDG reaction), 95 ^o^C for 5 min (abasic site cleavage) and 40 cycles of denaturation at 95 °C for 10 sec and annealing and extension at 60 °C for 30 sec. Primers and probe sets used to amplify viral DNA are indicated under the specific experiments in the results section. Primers and probe sets targeting RPP30 were used to calculate Frac U^DNA^ using the ΔΔC_t_ method.

**RT-qPCR of extracellular viral RNA.** Culture supernatants were collected from infected MC, MDM^EI^ or MDM^LI^ 7-days post infection unless otherwise specified. Supernatants were first spun to remove cellular debris, filtered using 0.22 μm filter and frozen at -80 ^0^C until use. RNA was extracted from 140 μl of culture soup using the Qiagen mini-Viral Prep kit according to the manufacturer's protocol. RNA (~3 µg) was treated with 2 units turbo-DNase (Invitrogen) in 1X turbo-DNase buffer for 30 minutes at 37 °C, to remove DNA carryover and then purified using Qiagen RNEasy kit. About 1.3 μg of RNA was used to generated cDNA with the Qiagen OmniScript cDNA preparation kit. Reverse transcription program was run at 37 ^o^C for an hour and 10 min at 95 ^o^C inactivation step according to the manufacturers protocol. Using the cDNA as input material and the Rotor Gene qPCR probe kit (Qiagen), the genomic HIV RNA copies in the supernatants of infected MC and MDM cultures were measured relative to a standard curve developed with the J-lat HIV integration standard cell line (see above). Thermal cycling conditions for qPCR consisted of 95 °C for 5 min, and 40 cycles of denaturation at 95 °C for 10 sec and annealing and extension at 60 °C for 30 sec. The measurements are reported as RNA copies per provirus present.

**Activity of HIV reverse transcriptase with dUTP and dTTP substrates.** To test whether or not a significant amount of uracil can be incorporated into reverse transcripts by HIV RTase we measured the enzymes capacity to distinguish between dTTP and dUTP incorporation using the SNE assay described above. For this we assembled two 100 μL reactions, with dNTP (1200 fmol) mixtures containing dTTP or dUTP respectively and with 600 fmol DNA probe. To these reactions 1 μL (2 units) of recombinant RTase (see above) was added, rapidly mixed and incubated at 37 C. Ten μL of the reactions were taken out at select timepoints (0, 10, 20, 30, 60, 120, 240, 2400 s), quenched in 40 μL of 98% formamide, 20 mM EDTA (pH 8.0), and resolved on a 15% denaturing polyacrylamide gel. The initial linear rates of dUTP or dTTP incorporation into the primer strand were measured by fitting data points at < 40% reaction.

**Sequencing of single viral reverse transcripts and genomic RNA copies.** This experiment was conducted in a series of steps that involves, preparation of cDNA and determination of a solution that contains a single cDNA copy/ 5ul of solution followed by nested PCR on a single clone and Sangar sequencing. First, monocytes and MDMs were infected with HIV-1^BaL^ as described above and culture supernatants were collected at day seven post infection for each cell type (MC and MDM), filtered and frozen at -80 °C until use. Extracellular viral RNA was extracted from 140 μl of culture supernatants using QIAamp Viral RNA mini kit (Qiagen), treated with turbo-DNase (Invitrogen) and re-purified using RNeasy Mini kit (Qiagen). cDNA was prepared using Omniscript cDNA preparation kit (Qiagen) and specific primers (ES7 & ES8, see table S1) targeting the V3/V4 region of the envelope gene of HIV. In order to aquire a cDNA solution that contains a single copy of HIV envelope gene, a serially diluted cDNA was quantiffied by qPCR using standard curve generated from the J-lat HIV integration standard cell line (see above) and by setting the highest cDNA dilution at 10 HIV copies per 5 µL of input cDNA solution which is the lowest detection limit of our assay. qPCR was done using Env200 forward and reverse primer sets (**Table S1**) and Rotor-gene SyBr green qPCR kit (Qiagen), in a final reaction volume of 25 µL (20 µL of master mix and 5 µL of input cDNA). Once a concentraion of 2 copies/µL of cDNA was achieved and confirmed by qPCR, it was further 2-fold serially diluted and subjected to nested PCR in five replicates to identify a dilution that contains a single clone per 5 µL of solution. Clonality of the candidate dilution was tested by a qPCR step using Env200 primer sets menationed above. A sample is deemed clonal if one replicate out of five tested positive according to Poisson statistics (1,2).

Once a solution that contains a single clone per 5 µL of solution is identified, several nested PCR reactions were performed to identify clones for sanger sequencing. The first PCR was performed using Platinum taq high-fidelity polymerase for 15 cycles in 5 replicate dilutions. PCR product was diluted 20x and a positive clone was identified by performing qPCR analysis on five replicate dilutions using primers targeting the amplified envelope fragment (Env200-F and Env200-R forward and reverse primers, **Table S1**) using Rotor Gene SyBr green kit (Qiagen) detection (3). The sample that gave a positive qPCR signal was then amplified by a second inner PCR with Platinum taq high-fidelity polymerase (Invitrogen) and E592 Forward and reverse primer sets (**Table S1**)(1). The thermocycler programs for the first and second PCR programs include initial denaturation at 94 ^o^C for 2 min, 15 cycles for first PCR or 35 cycles for second PCR consisting of 15 s of denaturation at 94 ^o^C, 15 s of annealing at 55 ^o^C and 1 min of extension at 68 ^o^C. The final extension was 2 min at 68 ^o^C and the samples were held at 4 ^o^C until further processing. Primers for the Env region are listed in **Supplemental Table S1**. The PCR products derived from amplifying the clonal isolates were run on a 1% agarose gel and the DNA bands were excised and purified with QIAamp gel purification kit (Qiagen). Purified DNA was directly sequenced by Sanger method with the forward and reverse primers listed in **Supplemental Table S1**.

Limiting dilution sequencing of proviral DNA was done in a similar fashion except that the input material for limiting dilution was genomic DNA. All clonal sequencing chromatograms were edited manually and aligned to our laboratory reference sequence of HIV-1 HXB3/BaL using ClustalW program in MEGA software v7.0.26 (MEGA). The bulk reference sequence for the envelope variable regions V3 and V4 (~550 bp) was determined from sequencing the population of proviruses and genomic RNAs obtained from infection of MOLT-4/CCR5 cells in our lab. The bulk sequences were unchanged after six generations of propagating HIV^BaL^ virus. The third and sixth generation viral stocks were used for all infections in this study.

**Over-expression of hUNG in MDMs**. Doxycycline inducible lentiviral particle (pCW57.1.FL.hUNG) was first generated by transfecting HEK 293 T cells with; 90 ug of pCW 57.1 hUNG, 90 ug of pMDLg/pRRE packaging plasmid (containing Gag & Pol, Addgene), 36 ug of pRSV-Rev packaging plasmid (containing Rev, Addgene), 18 ug of pMD2.g packaging plasmid (containing VSV-G envelope, Addgene). Transfection was performed using Lipofectamin 2000 (Invitrogen) following the manufacturers protocol. Lentivral particles were then concentrated over 20% sucrose cushion and ultracentrifugation at 28 k RPM for two hours. Viral titer was determined by the ELISA p24 antigen assay (Lenti-X p24 Rapid Titer Kit, TaKaRa). MDMs were then infected with this lentiviral particle at MOMI of 5 (0.1 pg p24/cell) in 24 well plate. Transduction was done by layering 200 µl of the LV on to the cells and incubating at 37 ^0^C over water bath with gentle rocking. 4 hours later 200 µl fresh medium was added to the cells and 24 hours later fresh media was changed. Induction was done by adding doxycycline into the culture medium to a final concentration of 1 µg/ml. Expression was determined by gel-based UNG activity assay.

**Gel based hUNG activity assay.** MDM were transduced with a lentiviral construct containing full length hUNG (pCW.57.1.FL.UNG) at MOI of 5 (0.1 pg p24/cell). Total cell extract was prepared at days 1 and 3 using cell lytic M (Sigma, C2978) according to the manufacturer’s instructions. Concentrations of cell extracts were determined by Bradford assay. hUNG activity of cell extracts was determined by gel-based UNG activity assay. Briefly, A 5’-FAM labeled 19-mer duplex (2.5 uM) containing a single dUMP on one strand was incubated with 10 µg of cell extract for one hour in 1x buffer (100 mM HEPES, 5 mM EDTA, 0.01% Brij-35, 5 mM DTT, PH 7.5) at room temperature. The reaction was stopped and quenched by addition of NaOH (20 mM final concentration) and heated for 30 minutes at 90^0^ C to cleave the abasic DNA product of the uNG reaction. One volume of 95% formamide loading buffer was added to the reaction which was electrophoresed through a denaturing 15% polyacrylamide gel containing 8 M urea for 40 minutes to resolve the 19 mer substrate from the 9 mer uracil excision product. The gel was then imaged on a Typhoon fluorescence imager.
