## Supplemental Table S1 for "Deficient uracil base excision repair leads to persistent dUMP in HIV proviruses during infection of monocytes and macrophages"

**S1 Table.** Primer and molecular beacon probe sequences for viral DNA sequences and RT-PCR measurements of UBER enzyme mRNA expression (5’🡪3’)

| **Target** | **Primer/probe sequences (5’-3’)** |
| --- | --- |
| Early RT | Forward: GCT AAC TAG GGA ACC CAC TGC TT |
| Early RT | Reverse: CAA CAG ACG GGC ACA CAC TGC TT |
| Early RT | Probe: FAM-AGC CTC AAT AAA GCT TGC CTT GAG TGC TTC-BHQ2 |
| Late RT (LRT) | Forward: TGTGTGCCCGTCTGTTGTGT |
| Late RT (LRT) | Reverse: GAGTCCTGCGTCGAGAGATC |
| Late RT (LRT) | Probe: FAM-CAGTGGCGCCCGAACAGGGA-BHQ2 |
| RPP30 | Forward: GATTTGGACCTGCGAGCG |
| RPP30 | Reverse: GCGGCTGTCTCCACAAGT |
| RPP30 | Probe: VIC-CTGACCTGAAGGCTCT-MGBNFQ |
| Alu | Forward: GCCTCCCAAAGTGCTGGGATTACAG |
| gag | Forward: CATGTTTTCAGCATTATCAGAAGGA |
| gag | Reverse: TGCTTGATGTCCCCCCACT |
| gag | Probe: FAM-CCA CCC CAC AAG ATT TAA ACA CCA TGC TAA-BHQ2 |
| Envelope (ES7) | Forward: CTGTTAAATGGCAGTCTAGC |
| Envelope (ES8) | Reverse: CACTTCTCCAATTGTCCCTCA |
| E592 | Forward: TGCCAATTTCACAGACAATGCT |
| E592 | Reverse: TCCAGGTCTGAAGATCTCGGA |
| Envelope 200 (Env200-F) | Forward: AGCACATTGTAACCTTAGAGCA |
| Envelope 200 (Env200-R) | Reverse: TGCATGGGAGTGTGATTGTGT |
| UNG^a^ | Forward: GCC AGA AGA CGC TCT ACT CC |
|  | Reverse: TCG CTT CCT GGC GGG |
| APE1^b^ | Forward: TGG AAT GTG GAT GGG CTT CGA GCC |
|  | Reverse: AAG GAG CTG ACC AGT ATT GAT GA |
| Pol β^b^ | Forward: GGC AGT TTC AGA GGT GC |
|  | Reverse: GGC AAA CAC CCA TGA ACT TT |
| Lig III^b^ | Forward: GAT CAC GTG CCA CCT ACC TTG T |
|  | Reverse: GGC ATA GTC CAC ACA GAA CCG T |
| SAMHD1^c^ | Forward: GGA TTA CTA AAA ACC AGG TTT CAC AAC T |
|  | Reverse: TGT CGT TCC ATT CCT TTT TTT GA |
| A3A^d^ | Forward: GAA GGG ACA AGC ACA TGG AAG C |
|  | Reverse: ATC TAC TTG ATC GGG AGC ATA C |
| A3G^d^ | Forward: GGT GTA TTC CGA ACT TAA GTA C |
|  | Reverse: CAA GGA AAC CGT GTT TAT GTG G |
| 18S^a^ | Forward: TGT GCC GCT AGA GGT GAA ATT |
|  | Reverse: TGG CAA ATG CTT TCG CTT T |

^a^Virology Journal (2012) *9*, 230; ^b^[Mol Cell Biochem.](https://www.ncbi.nlm.nih.gov/pubmed/28887667) (2018) *441*, 201; ^c^J. Biol. Chem. (2013) *288*, 9284; ^d^J Invest Dermatol. (2017) *137*, 810.
