## Supplemental Figure 2 for "Deficient uracil base excision repair leads to persistent dUMP in HIV proviruses during infection of monocytes and macrophages"

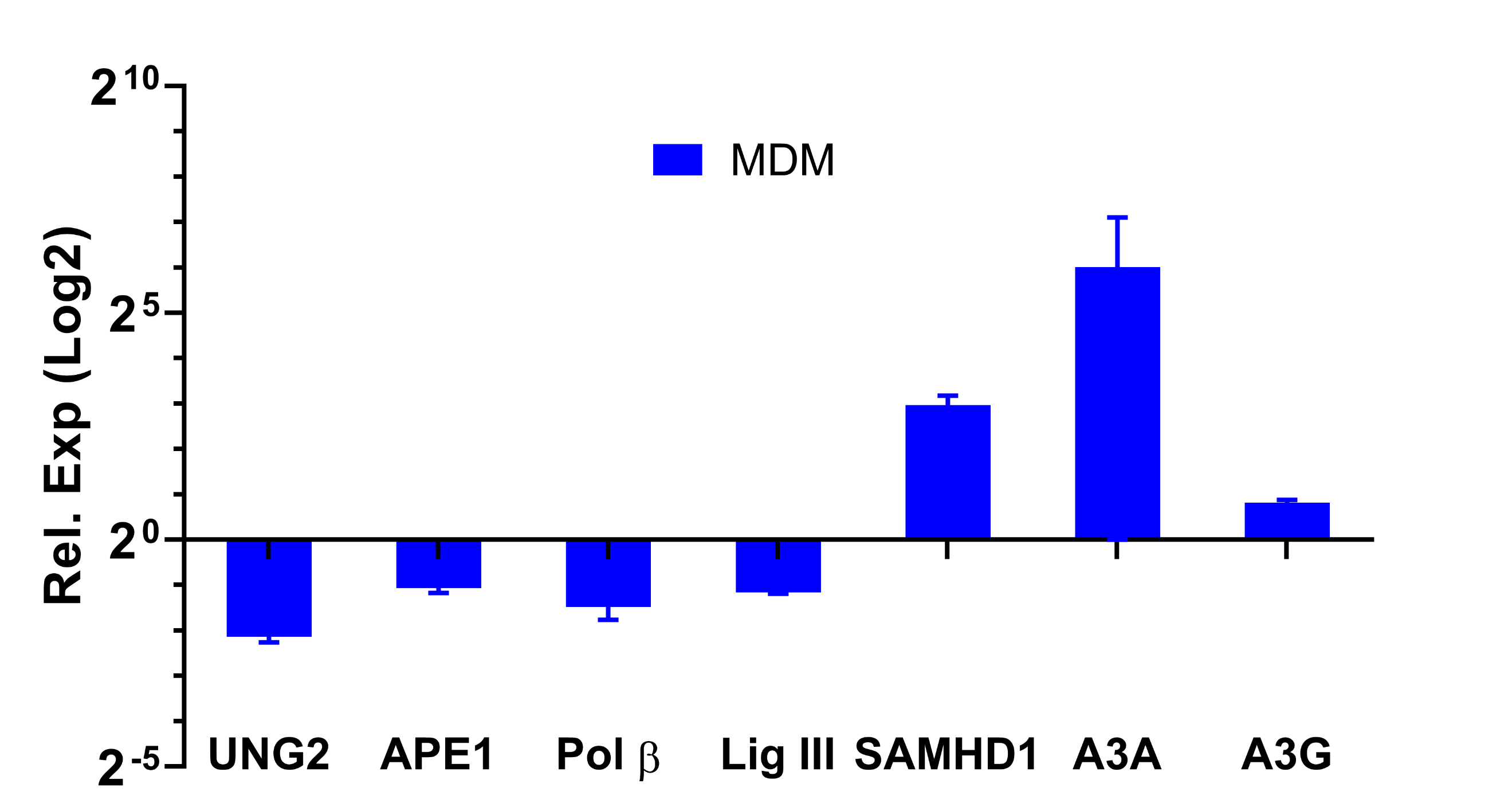

**S2 Fig.** **Determination of UBER mRNA expression levels in MDMs and comparison with the HAP1 dividing cell line.** Relative mRNA expression levels of UBER enzymes in MDM relative to HAP1 dividing cells. Total RNA was extracted from MDMs after seven days differentiation from MC. All qPCR reactions were done in triplicate.
