## Supplemental Table S2 for "Deficient uracil base excision repair leads to persistent dUMP in HIV proviruses during infection of monocytes and macrophages"

**S2 Table.** Clonal mutation analysis of HIV proviral DNA isolated from infected MDM at 7 dpi

| **Seq**  **(+ strand)** | **Type** | **Position** | **Possible mechanism^a^** | **Amino Acid mutations** |
| --- | --- | --- | --- | --- |
| TGTACGAGACC | A-->G | 903 | i. (-) strand C mispairing with A, or ii. M-RA^b^ | E322G |
| AATACGAGAAA | A-->G | 921 |  | None |
| ATACAAGAAAAA | A-->G | 922 | (-) strand C mispairing with A | None |
| ACAAGGAAAAG | A-->G | 925 |  | None |
| GGAGGTATAA | A-->G | 983 |  | D326G |
| AGAGAGCAATT | A-->G | 1062 |  | None |
| CTTTAGGCACT | A-->G | 1091 |  | None |
| TTGTGGCGCAC | A-->G | 1123 |  | T374A |
| TCAACGCAACT | A-->G | 1171 |  | T399I |
| CAAATGTTACA | A-->G | 1340 |  | I450V |
| CTATTGACAAG | A-->G | 1356 |  | None |
| AAGACTCAACA | C-->T | 967 | (-) strand A mispairing with C | P299L |
| TGTTATTGAAG | C-->T | 1200 |  | None |
| TAACATTGTAG | C-->T | 1218 |  | T409I |
| ACACTTCCATG | C-->T | 1242 | i. (-) strand A mispairing with C ii. M-RA^b^ | None |
| TTAATCGTACA | T-->C | 898 | (-) strand G wobble pairing with U | C296R |
| AAAACGGAAT | T-->C | 1021 |  | W339R |
| ACAGTCTTAAT | T-->C | 1133 |  | F377L |
| GAAATGAATTG | T-->G | 895 | (-) C mispairing with U | None |
| GCATTGTATACA | T-->G | 957 |  | None |
| AGTTAGAAAAT | T-->G | 1049 |  | I348R |
| AGGCAAAGCAT | G-->A | 950 | i. (-) strand U wobble pairing with G ii. A3A cytosine deamination | None |
| GGCAGAAAGTA | G-->A | 1279 | i. (-) strand U wobble paring with G ii. A3G cytosine deamination | E430K |
| AACAACTTATA | A-->C | 1261 | (-) strand G mispairing with A | None |
| ATTATCAACAT | A-->C | 1266 |  | None |
| TGAATTAATCT | G-->T | 881 | (-) strand A mispairing with G | stop codon 290 |
| CTGAAAGAATC | T-->A | 880 | (-) strand U mispairing with U | N289K |

^a^Mutational events leading to base pair mismatches during (+) strand synthesis are not considered because no mismatches were ever detected in the sequencing studies. Mismatches occurring during (-) strand synthesis are more consistent with the absence of observable mismatches because the (+) strand RNA template is degraded before second strand synthesis, which removes the mismatch and results in a base pair substitution after reverse transcription. ^b^M-RA indicates a misalignment-realignment event during reverse transcriptase extension.
