## Supplemental Figure 3 for "Deficient uracil base excision repair leads to persistent dUMP in HIV proviruses during infection of monocytes and macrophages"

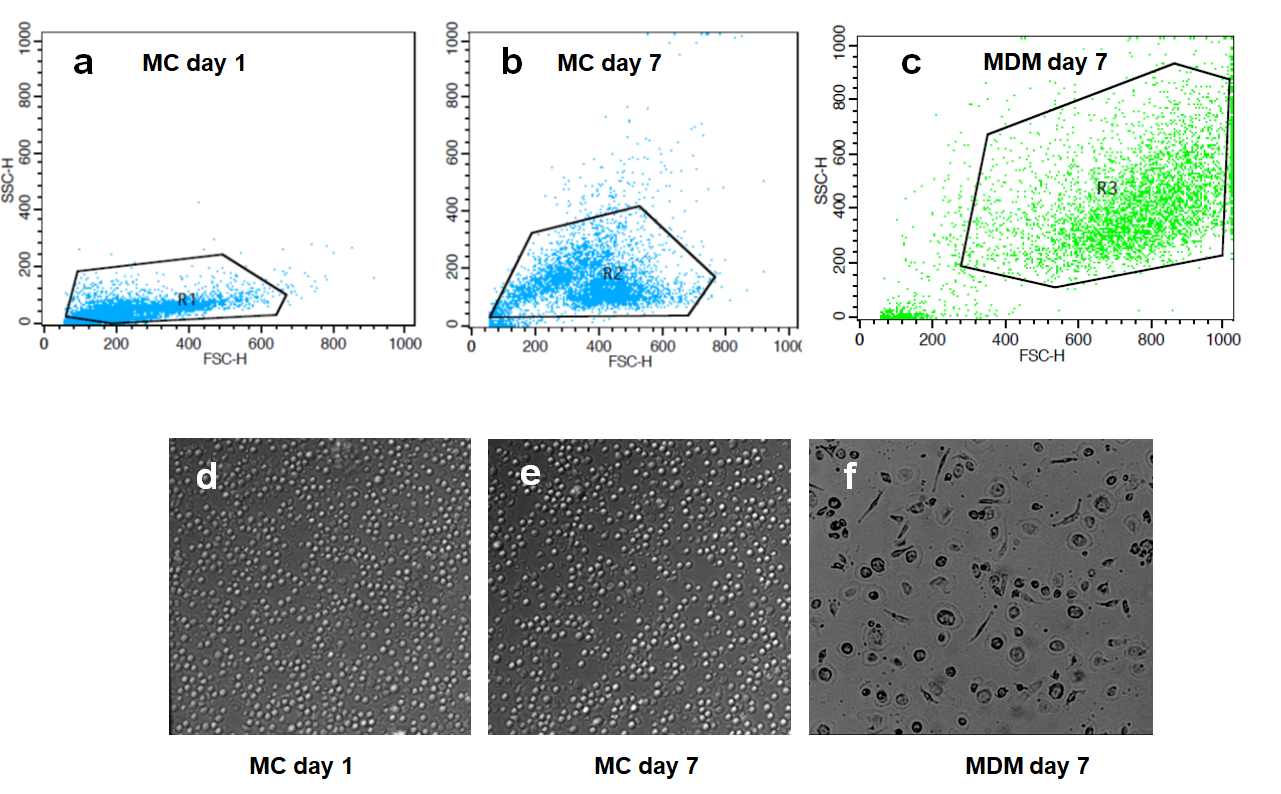

**S3 Fig. Morphological and granulation differences between MDMs and MCs. (a)** Flow cytometry analysis of monocytes immediately after purification. (**b**) Flow cytometry analysis of monocytes cultured in suspension under non-adherent conditions for seven days. (**c**) Flow cytometry analysis of fully differentiated MDM (cultured for 7 days under adherent conditions in the presence of M-CSF). (**d**) Light microscope image of monocytes immediately after purification (20x magnification, 5x zoom). **(e**) Light microscope image of monocytes after 7 days of culture under non-adherent conditions in the absence of M-CSF (20x magnification, 5x zoom). (**f**) Light microscope image MDM after 7 days of culture in the presence of M-CSF (20x magnification, 4x image reduction).
