## Supplemental Table S3 for "Deficient uracil base excision repair leads to persistent dUMP in HIV proviruses during infection of monocytes and macrophages"

**S3 Table.** Clonal mutation analysis of HIV proviral DNA isolated from infected MC at 7 dpi

| **Seq** | **Type** | **Position** | **Possible mechanism^a^** | **AA mutation** |
| --- | --- | --- | --- | --- |
| GAAAAGGTATA | A-->G | 930 | (-) strand C mispairing with A | S306R |
| ATAGGGGATAT | A-->G | 983 |  | None |
| GAATGGATCTG | A-->G | 883 | i. (-) strand C mispairing with A, or ii. M-RA^b^ | None |
| TAAAACCATAA | T-->C | 861 | (-) strand G wobble pairing with U | I283T |
| GAGCACTTTAT | T-->C | 957 |  | None |
| AAGCACTCCTC | T-->C | 1127 |  | E381G |
| AATTAAATGTT | G-->A | 1327 | i. (-) strand U wobble pairing with G ii. A3A cytosine deamination | R444K |
| TGGCAAGAAGT | G-->A | 1280 | i. (-) strand U wobble pairing with G, or ii. A3A cytosine deamination, or iii. M-RA^b^ | None |
| TAACATTGTAG | C-->T | 1219 | (-) strand A mispairing with C | T408I |
| AGTTAGAAAAT | T-->G | 1051 | (-) strand C mispairing with U | I347R |
| GCATTTTATAC | G-->T | 959 | (-) strand A mispairing with G | None |

^a^Mutational events leading to base pair mismatches during (+) strand synthesis are not considered because no mismatches were ever detected in the sequencing studies. Mismatches occurring during (-) strand synthesis are more consistent with the absence of observable mismatches because the (+) strand RNA template is degraded before second strand synthesis, which removes the mismatch and results in a base pair substitution after reverse transcription. ^b^M-RA indicates a misalignment-realignment event during reverse transcriptase extension.
