## Supplemental Figure 4 for "Deficient uracil base excision repair leads to persistent dUMP in HIV proviruses during infection of monocytes and macrophages"

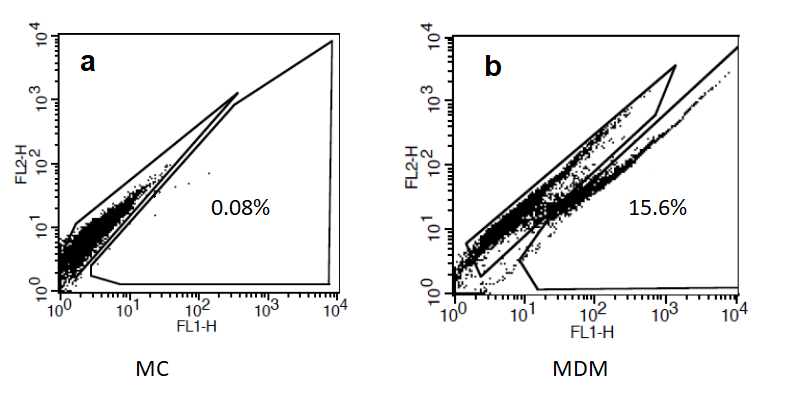

**S4 Fig. HIV-eGFP expression in MC and MDM.** (**a**) MC were infected with HIV^NL4-3(eGFP)^ immediately after isolation at an moi of ten. At 7-days post infection, eGFP expression was measured by flow cytometry. Even though GFP fluorescence is very low, viral reverse transcripts are abundant (main text). (**b**) Fully differentiated MDM were infected with HIV^NL4-3(eGFP)^ at an moi of ten. At 7-days post infection, eGFP expression was measured by flow cytometry.
