## Supplemental Table S4 for "Deficient uracil base excision repair leads to persistent dUMP in HIV proviruses during infection of monocytes and macrophages"

**S4 Table:** Clonal mutation analysis of dUMP-depleted HIV proviral DNA isolated from infected MDM at 7 dpi

| **Seq** | **Type** | **Position** | **Mechanism^a^** | **AA mutations** |
| --- | --- | --- | --- | --- |
| TTTATGCAACA | A-->G | 962 | (-) strand C mispairing with A | T319A |
| ACAACGGGAGA | A-->G | 966 |  | None |
| TACATGTAGGA | A-->G | 939 |  | I309V |
| CAACAGCAATA | A-->G | 916 |  | None |
| AGGAGGTATAA | A-->G | 985 |  | None |
| CAGAAGTTGTG | A-->G | 1119 |  | None |
| TCATCGAATAT | A-->G | 1335 |  | N449D |
| GGAATATTACT | G-->A | 1197 | i. (-) strand U wobble pairing with G ii. A3A cytosine deamination | V396I |
| TAGATATTCAT | G-->A | 1330 |  | C445Y |
| GGAATAACACT | G-->A | 1028 |  | D341N |
| TAAGAAAACAA | G-->A | 1060 |  | E352K |
| CTGAAAAGTCA | G-->A | 1205 |  | None |
| CCCAATAACAA | C-->T | 912 | (-) strand A mispairing with C | None |
| TGACATTTTAA | C-->T | 1032 |  | T342I |
| GGACCTAGAAA | C-->T | 1112 |  | L317S |
| AGCAACGTATG | T-->C | 1295 | (-) strand G wobble pairing with U | None |
| CAAAACGGAAT | T-->C | 1023 |  | W338R |
| TCCCACGCAGA | T-->C | 1248 |  | C418R |
| GCAGATCATTG | G-->T | 954 | (-) strand A mispairing with G | L318F |

^a^Mutational events leading to base pair mismatches during (+) strand synthesis are not considered because no mismatches were ever detected in the sequencing studies. Mismatches occurring during (-) strand synthesis are more consistent with the absence of observable mismatches because the (+) strand RNA template is degraded before second strand synthesis, which removes the mismatch and results in a base pair substitution after reverse transcription.
