## Supplemental Figure 5 for "Deficient uracil base excision repair leads to persistent dUMP in HIV proviruses during infection of monocytes and macrophages"

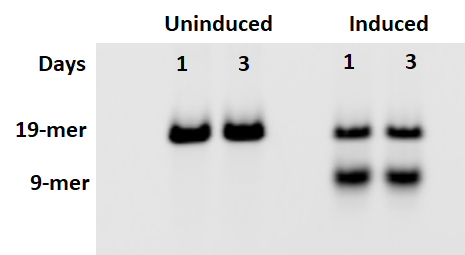

**S5 Fig. Activity of lentiviral transduced hUNG2 in MDM cell extracts under uninduced and induced conditions.** MDM were transduced with a lentiviral construct containing full length hUNG2 (pCW.57.1.FL.UNG) at MOI of 5 (0.1 pg p24/cell). Total cell extracts were prepared at days 1 and 3 post-transduction and protein concentrations were determined by the Bradford assay. hUNG activity in cell extracts was determined by gel-based UNG activity assay using equal amounts of total protein and a 19-mer uracil-containing ssDNA substrate with a FAM label on the 5′-end (see Methods). Excision of uracil results in a 5′-FAM labeled 9mer product band.
