## Supplemental Table S5 for "Deficient uracil base excision repair leads to persistent dUMP in HIV proviruses during infection of monocytes and macrophages"

**S5 Table.** Clonal mutation analysis of extra cellular viral RNA extracted from infected MDM culture supernatants at 7 dpi^a^

| **Sequence** | **Type** | **Env position** | **Coding changes** |
| --- | --- | --- | --- |
| AAATAGGATAG | A-->G | 1040 | None |
| TTGGGGATAAA | A-->G | 1073 | N357D |
| CCCAGGAATTG | A-->G | 1116 | E370G |
| TTTTGATTGT | A-->G | 1136 | N378D |
| AGTCAGATAAC | A-->G | 1211 | N407D |
| GAGCAGTGTAT | A-->G | 1294 | M435V |
| CCATCGGAGGA | A-->G | 1312 | R436G |
| GAGTCGAATAA | A-->G | 1209 | None |
| TTAATCGTACA | T-->C | 899 | C296R |
| GAGCACTTTAT | T-->C | 996 | None |
| GAGCATCTTATA | T-->C | 997 | F317P |
| TAAGATAAGCA | C-->T | 991 | Nonsense |
| AATAATACTGT | C-->T | 1215 | None |
| GTAATACAACA | T-->A | 1165 | None |
| GGTCCAGAGGA | T-->A | 1375 | None |
| GAAAAGAACAC | T-->G | 1227 | N412K |
| GTATAAATATA | C-->A | 935 | H308N |
| ^a^Possible mutational mechanisms are not indicated because of the ambiguity in assigning mutations to reverse transcription or RNA pol II transcription. | | | |
