## Supplemental Figure 6 for "Deficient uracil base excision repair leads to persistent dUMP in HIV proviruses during infection of monocytes and macrophages"

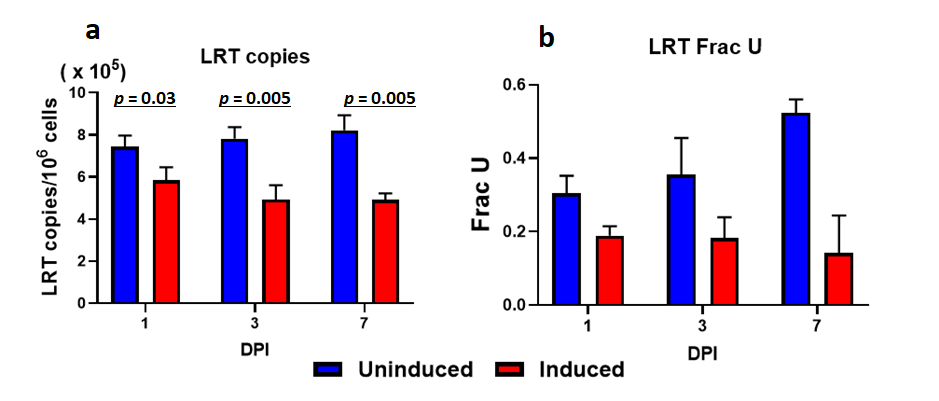

**S6 Fig. Effect of over expression of hUNG2 in MDMs on total HIV DNA.** Fully differentiated MDM were first transduced with inducible lentiviral construct expressing full length hUNG at MOI of 5 (0.1 pg p24/cell) and 3 days later induced with doxycycline (1ug/ml). 1-day after induction, MDM were then infected with HIV^NL4-3^ single round virus at MOI of 0.5 (0.05 pg p24/cell). Total DNA was extracted at days 1, 3 and 7 and (**a**) LRT copies and (**b**) Frac U were measured by Ex-qPCR.
